# Synaptic Encoding of Vestibular Sensation Regulates Movement Timing and Coordination

**DOI:** 10.1101/2021.07.05.451142

**Authors:** Kyla R. Hamling, Katherine Harmon, Marie Greaney, Zoë Dobler, Yukiko Kimura, Shin-ichi Higashijima, David Schoppik

## Abstract

Vertebrate vestibular circuits use sensory signals derived from the inner ear to guide both corrective and volitional movements. A major challenge in the neuroscience of balance is to link the synaptic and cellular substrates that encode body tilts to specific behaviors that stabilize posture and enable efficient locomotion. Here we address this problem by measuring the development, synaptic architecture, and behavioral contributions of vestibulospinal neurons in the larval zebrafish. First, we find that vestibulospinal neurons are born and are functionally mature before larvae swim freely, allowing them to act as a substrate for postural regulation. Next, we map the synaptic inputs to vestibulospinal neurons that allow them to encode posture. Further, we find that this synaptic architecture allows them to respond to linear acceleration in a directionally-tuned and utricle-dependent manner; they are thus poised to guide corrective movements. After loss of vestibulospinal neurons, larvae adopted eccentric postures with disrupted movement timing and weaker corrective kinematics. We used a generative model of swimming to demonstrate that together these disruptions can account for the increased postural variability. Finally, we observed that lesions disrupt vestibular-dependent coordination between the fins and trunk during vertical swimming, linking vestibulospinal neurons to navigation. We conclude that vestibulospinal neurons turn synaptic representations of body tilt into defined corrective behaviors and coordinated movements. As the need for stable locomotion is common and the vestibulospinal circuit is highly conserved our findings reveal general mechanisms for neuronal control of balance.

## Introduction

To remain balanced during locomotion, animals actively modulate the timing and strength of their trunk and limb/effector movements [1–5]. These computations are performed by brainstem neurons that integrate synaptic inputs from the inner ear with extravestibular information to transform sensed imbalance into specific behaviors. Understanding this transformation requires both mapping the underlying synaptic organization and defining the specific contribution of vestibular neurons to behavior. Vestibulospinal neurons are descending projection neurons conserved across vertebrates that are well-poised to regulate balance [6–10]. They have somata in the lateral vestibular nucleus [11, 12] and receive convergent excitatory [13, 14] and inhibitory vestibular input [15, 16] as well as a wide variety of extravestibular inputs [17].

Vestibulospinal neuron development and their precise contribution to behavior remain active areas of research, largely limited by the complexity of both the mammalian brain and tetrapod locomotion. In decerebrate cats, vestibulospinal neurons encode body tilts, are active during extensor muscle activation during locomotion [14, 18–20], and relay VIII^th^ nerve activity to ipsilateral extensor muscles [21, 22]. Recent technological improvements permit targeting of genetically-defined populations of vestibulospinal neurons in mice. These tools have been used to illuminate how connectivity to downstream areas develops [23–26] and to further characterize vestibulospinal neuron-dependent hindlimb responses following imposed instability [27, 28]. Taken together, while the vestibulospinal circuit encodes instability and influences muscle tone, its contribution to postural behaviors remains challenging to discern.

Recent work has used larval zebrafish to explore vertebrate vestibular circuits, but our understanding of neuronal development and synaptic architecture remains incomplete. In larval zebrafish, spinal-projecting hindbrain neurons [29–31] encode postural information from the utricle [32]. However vestibulospinal development remains poorly understood, making links to behavior tenuous. Elegant work used intracellular recording to characterize convergent excitatory otolithic inputs to vestibulospinal neurons [32], and imaging to map brain-wide responses to imposed destabilization [33, 34]. However this work focused primarily on excitatory drive from the utricle. Though key experiments remain, the larval zebrafish is thus well-established as a means to explore the neural basis of vertebrate postural control.

Swimming offers a particularly tractable means to assay neuronal contributions to balance computations. The biophysical and biomechanical contributions to swimming are straightforward to delineate [35, 36], allowing dissociation of passive and active (i.e. neuronal) contributions. For example, larval zebrafish swim with short discontinuous propulsive movements called bouts. Zebrafish maintain their preferred near-horizontal pitch using two computations: they learn to time these bouts to occur when pitched off-balance and they counter-rotate their bodies during bouts [37, 38]. Loss-of-function experiments have linked vestibular sensations to coordination of fin and body movements during bouts during vertical navigation [39], and spinal contractions for postural righting [40]. Swim behavior is therefore an excellent means to understand the specific contributions of particular neurons to postural control.

Here, we assayed the development, synaptic, circuit, and behavioral properties of larval zebrafish vestibulospinal neurons. We found that vestibulospinal neurons are all born well before larvae begin to swim freely. Consistent with their early birth date, *in vivo* whole-cell recordings showed that vestibulospinal neurons were electrophysiologically mature, poising them to participate in behavior. We used voltage-clamp recordings together with inner-ear lesions to map the organization of synaptic inputs to these neurons, delineating both vestibular and extra-vestibular contributions. Vestibulospinal neurons responded reliably in a directionally-specific manner to body displacement (linear acceleration), allowing them to relay perceived instability. To determine their specific contribution to behavior, we photoablated neurons and monitored posture and locomotion in freely swimming fish. Without vestibulospinal neurons, posture was more variable. We used a model of swimming to determine that disrupted movement timing and corrective kinematics could together account for increased postural variability. Finally, we found that fin/body coordination was impaired after neuronal loss. Taken together, our work (i) defines the development and synaptic architecture of vestibulospinal neurons and (ii) determines their specific contribution to computations that stabilize posture and facilitate vertical navigation.

## Results

### Vestibulospinal neurons are an anatomically-identifiable population born well before spontaneous locomotion

In mammals, spinal-projecting neurons innervated by the vestibular branches of the VIIIth cranial nerve are located in anatomically and cytoarchitectonically distinct hindbrain regions termed the lateral vestibular nucleus [41]. We optically (i.e. non-invasively) labeled all spinal projecting neurons in the larval zebrafish to allow a more comprehensive characterization of putatively homologous neurons relative to previous work [29]. We observed a distinct cluster of spinal-projecting neurons in the lateral region of hindbrain rhombomere 4 near the VIII^th^ cranial nerve and the lateral dendrite of the Mauthner cell [29, 30] (Figures 1A, 2A). Clusters contained 27±5 neurons per hemisphere (n=6 hemispheres, 3 fish), distributed evenly dorsal and ventral to the Mauthner lateral dendrite (Figure 1B, 15±5 above, 12±3 below, paired t-test p=0.32). Neurons with large cell bodies, presumably homologous to Deiter’s neurons [42], were located exclusively in the ventral region while smaller neurons were found dorsally. Based on their location, projections, and anatomy we classified these neurons as vestibulospinal.

**Figure 1:**
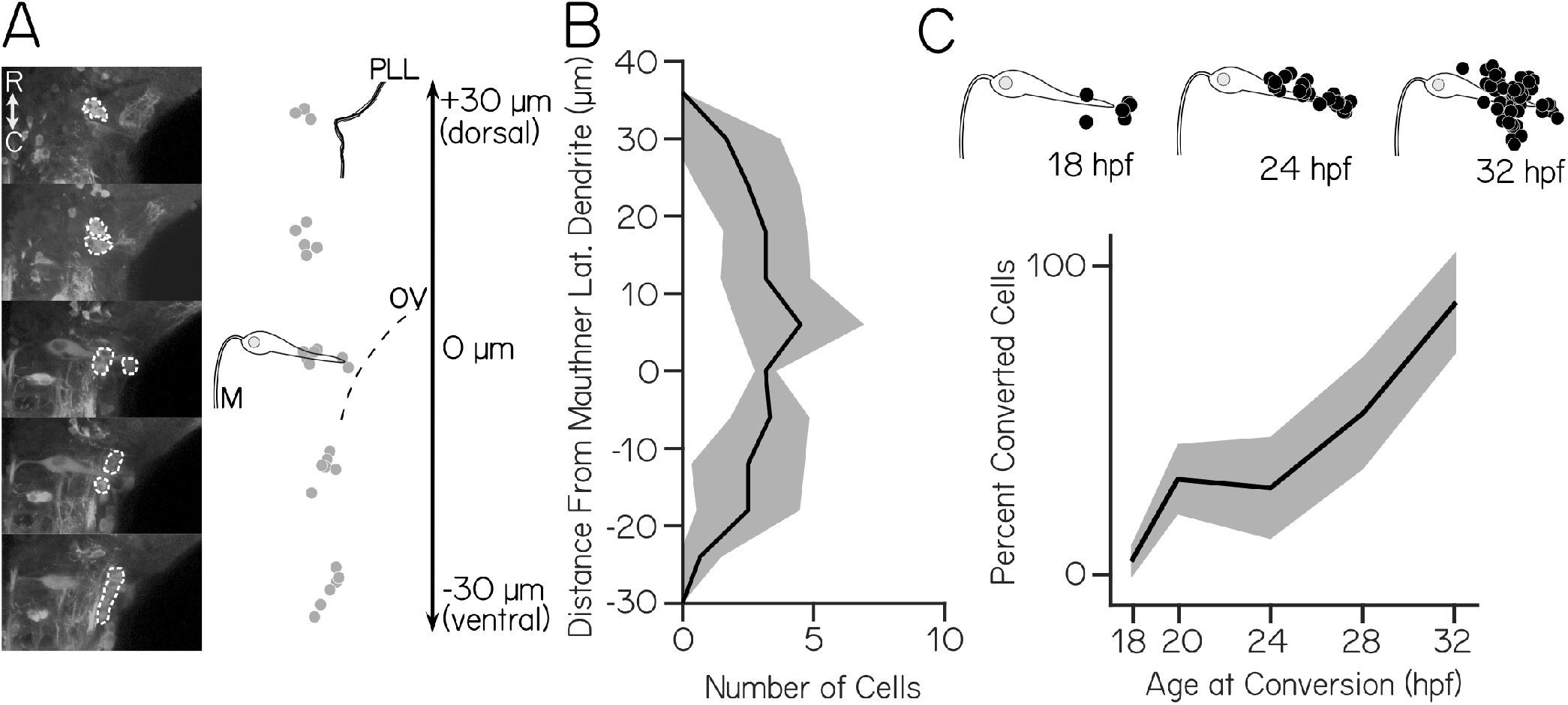
Vestibulospinal neurons are born well before larvae swim or stabilize posture. (A) Left: Images of optically backfilled spinal projecting neurons (dashed white) in five dorsoventral planes covering ± 30 um around the Mauthner (M) lateral dendrite, somata. Right: representative neuron location (gray circles) in one hemisphere and the Mauthner lateral dendrite (B) Mean dorsoventral distribution of neurons relative to the Mauthner lateral dendrite per hemisphere (± 1 S.D.) (n=6 fish) (C) Top: neurons (black circles) born at 18, 24, and 32 hpf relative to the Mauthner cell. Bottom: Percent of vestibulospinal neurons born increases steadily to near 100% between 18 and 32 hpf (mean ± 1 S.D.)

**Figure 2:**
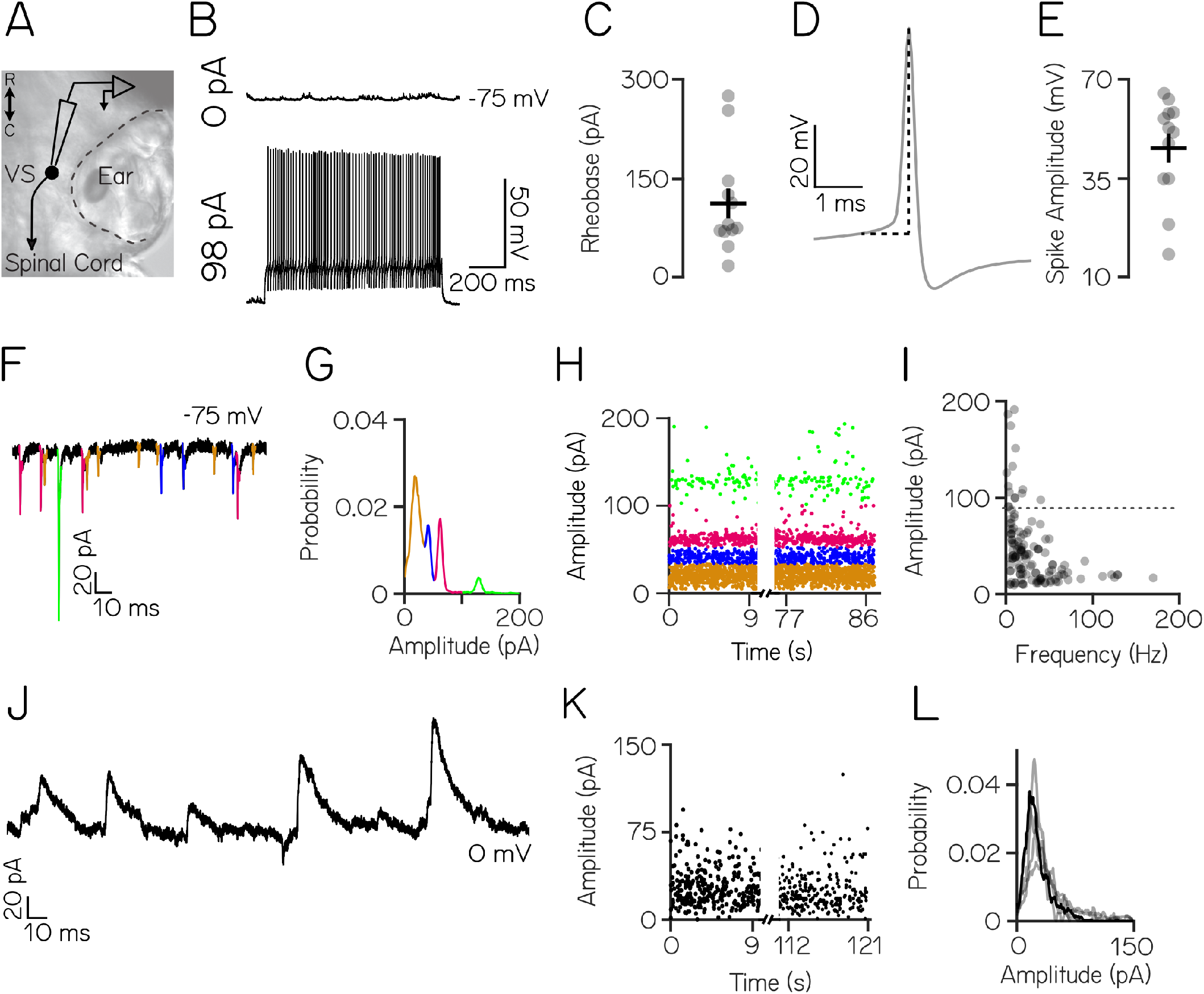
Larval zebrafish vestibulospinal neurons display mature intrinsic and synaptic properties. (A) Schematic of spinal-projecting vestibulospinal (VS) neuron targeted for electrophysiology (B) Vestibulospinal response to current injection (98 pA) (C) Rheobase (±SEM) across 21 cells (D) Example action potential waveform with amplitude (dotted line) (E) Action potential average amplitude (±SEM) (F) Distinct amplitudes (color) in excitatory post-synaptic spontaneous currents (EPSCs) from a neuron held at −75mV (G) EPSC amplitudes from a single vestibulospinal cell show 4 distinct probability peaks, or “bins” (color) (H) EPSC bins are stationary in time (I) EPSC amplitude as a function of frequency for all bins in all vestibulospinal cells (116 bins, from 36 cells), dashed line differentiates the top 5% (“high amplitude”) of bins (J) Representative current trace from a vestibulospinal neuron held at 0 mV (K) Inhibitory post-synaptic current (IPSC) amplitudes over time (L) IPSC amplitudes for the example neuron in J/K (black line) and other neurons (gray lines) do not show multiple peaks (n=5)

To determine when neurons in the vestibulospinal cluster develop we used photoconversion of the photolabile protein, Kaede, to define their time of terminal differentiation, or “birthdate,” [43, 44]. The earliest-born neurons were present at 18 hours postfertilization (hpf) (4±5%) and most were born by 32 hpf (88±17%), with the majority (60%) born between 24-32 hpf (Figure 1C). Larval zebrafish can produce postural righting behaviors to orient themselves dorsal-up as early as 3 days post-fertilization [40, 45]. These data indicate that vestibulospinal neurons are present before the onset of normal postural behaviors.

### Vestibulospinal neurons have low spontaneous firing rates but receive strong excitatory and inhibitory synaptic input

In other species, mature vestibulospinal neurons integrate synaptic input and fire action potentials to relay sensed imbalance to corrective effectors. We first performed *in vivo* whole cell patch clamp recordings in the dark in 4-12 dpf fish (n=21 cells). Dye in the recording solution allowed *post-hoc* assignation of vestibulospinal identify by visualization of descending axons. Neurons had high input resistance (236±130 MΩ) and resting membrane potential of −67±5 mV (Table 1). At rest, approximately half (12/21) of vestibulospinal neurons showed no spontaneous action potentials (Figure 2B, top); the remaining 9 had an average firing rate at rest of 3.2±5.3 Hz. To determine if the absence of spontaneous activity reflected an immature phenotype we delivered steps of depolarizing current. Silent vestibulospinal cells could sustain a high tonic firing rate (90.3±68.5 Hz) following current injection (largest depolarizing step per cell: 84-322 pA) (Figure 2B, bottom). The high rheobase of 111.8±78.9 pA (Figure 2C), suggests the low spontaneous rate reflects a high spiking threshold. Evoked action potentials had a mature waveform with a mean spike amplitude of 48.2 mV (Figure 2D,E). Neither variation in spike amplitude nor rheobase was correlated with age (Spearman’s correlation; p=0.17 spike amplitude, p=0.97 rheobase), suggesting that by the time fish swim and stabilize posture, vestibulospinal neurons, even those silent at rest, have developed mature characteristics.

**Table 1:**
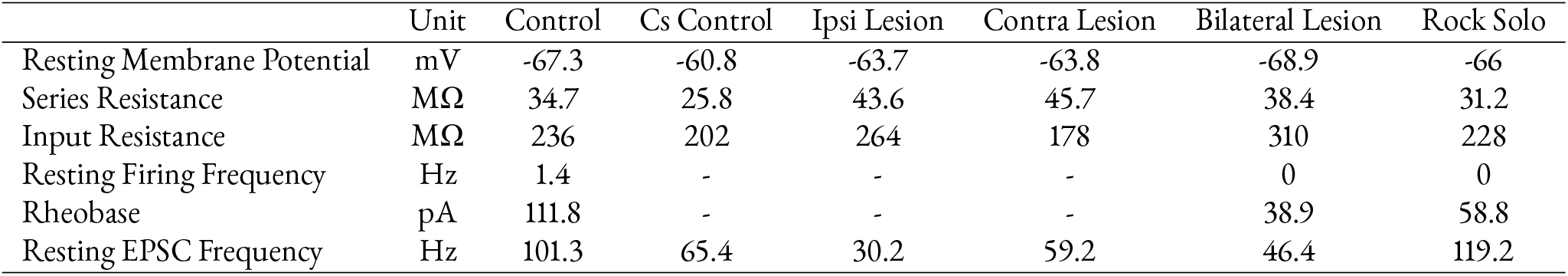
Spontaneous properties across conditions

The observation that vestibulospinal neurons have a low/absent resting firing rate might reflect the absence of strong synaptic input at rest. To test this possibility we recorded excitatory post-synaptic currents (EPSCs) in voltage clamp. Neurons received dense synaptic EPSCs (Figure 2F) with a mean EPSC frequency of 103.2±54.0 Hz (n=36 neurons). EPSCs showed a wide range of amplitudes (median range 136.0 pA). Amplitude distributions were multimodal in almost all cells (97%), with distinct peaks in the probability distribution of EPSC amplitudes (Figure 2G). To characterize these peaks, we assigned EPSCs to amplitude ranges that encompassed each peak in the distribution (Figure 2G, line colors). These bins remained stable over time (Figure 2H). Neurons had a mean of 3±1 distinct bins (116 bins from 36 cells), with a median event amplitude per bin of 39.0±27.0 pA and a median event frequency per bin of 19.4±21.7 Hz. Bin amplitude and frequency were inversely related (Figure 2I). Consistent with birthdating and intrinsic properties, neither bin amplitude nor frequency were correlated with age, suggesting mature excitatory inputs to vestibulospinal neurons (Pearson correlation; p=0.69 bin amplitude, p=0.47 bin frequency).

Distinct EPSC bins might reflect input from single VIII^th^ nerve afferents with different stable resting amplitudes [46, 47] as proposed for comparable recordings done in the light [32]. To test if EPSCs within a distinct amplitude bin in our recordings derived from a single afferent neuron, we compared empirical within-bin refractory period violations to a model estimate of expected violations accounting for noisy bin assignations and within-bin event frequency (Figure S1). The majority (98/116) of amplitude bins had more violations than expected, suggesting that EPSCs that fall within the same amplitude bin likely originate from multiple afferent neurons. Bins that passed our refractory test, indicating a putatively unitary afferent origin, had higher amplitude and lower frequency EPSCs (median 54.9 pA vs. 31.1 pA, median 9.7 Hz v. 24.4 Hz). The inter-event intervals of the putative unitary bins were consistent with irregular input (coefficient of variation = 0.96±0.11). Some bins (31/116) exhibited higher than expected refractory period violations, which might reflect compound events (i.e. mixed electrical and chemical synapses). However, waveforms of high-probability compound events did not have the expected shape of a classical electrochemical synapse with a high amplitude electrical event followed by a stereotyped low-jitter chemical event (S1F-G). We conclude that in our recordings most bins are comprised of multiple inputs, but some high-amplitude, low-frequency event bins are consistent with single irregularly-firing VIII^th^ nerve afferents.

To assay inhibitory inputs at rest, we performed a separate set of voltage-clamp experiments to isolate inhibitory post-synaptic currents (IPSCs). Neurons received spontaneous IPSCs (Figure 2J) with a mean frequency of 22.4±7.4 Hz (range 13.8-32.5 Hz) and a mean amplitude of 29.1±7.3 pA (range 22.5-39.2 pA) (n=5 neurons). Unlike excitatory inputs, spontaneous inhibitory currents did not exhibit distinct event amplitude peaks (Figure 2K,L). We conclude that neurons receive dense excitatory and inhibitory input, and, when sufficiently depolarized, can fire sustained trains of action potentials.

### High amplitude excitatory inputs derive from the ipsilateral utricle

Electrophysiological characterization indicates that vestibulospinal neurons would be poised to encode and stabilize posture if sufficient synaptic input arrived during a body tilt. Previous inner-ear loss-of-function studies established that the utricle is the dominant source of sensory information about body tilts in larval zebrafish [48, 49]. We assayed if synaptic input to vestibulospinal neurons was of utricular origin by recording spontaneous synaptic activity in vestibulospinal neurons after inner-ear lesions either ipsilateral or contralateral to the recorded neuron (Figure 3A-B). First, the number of EPSC amplitude bins per cell differed across lesion conditions (Kruskal-Wallis H(2)=10.2, p=0.006) (Figure 3C). After ipsilateral lesion, neurons had fewer bins (median 1 vs 3; n=9 lesion, n=5 control; p=0.0084, Wilcoxon rank sum test). In contrast there was no change after contralateral lesion (median 3 bins; n=6; p=0.095, Wilcoxon rank sum test). Further, the amplitude of EPSC bins also differed across conditions (Kruskal-Wallis H(2)=11.7, p=0.003) (Figure 3D). EPSC bins after ipsilateral lesion were lower amplitude than controls (median 11.4 pA vs. 43.2 pA; p=0.0011, Wilcoxon rank sum test), but contralateral lesions did not affect EPSC bin amplitudes (median 25.4 pA; p=0.1753 Wilcoxon rank sum test). EPSC bin frequency was not changed across lesion conditions (Kruskal-Wallis H(2)=0.06, p=0.97) (Figure 3E). Finally, we observed that the frequency of inhibitory synaptic inputs in vestibulospinal neurons could be disrupted by both ipsilateral and contralateral lesions (Figure S2). We conclude that ipsilateral lesions selectively disrupt high-amplitude, low-frequency EPSCs (Figure 3F). In contrast, lower-amplitude EPSCs persist after both ipsilateral and contralateral lesions, which might reflect either an extra-vestibular origin or an incomplete lesion. Our data suggest that individual neurons receive: 1) high-amplitude excitatory inputs exclusively from the ipsilateral utricle, 2) utricle-independent low-amplitude excitatory inputs, 3) inhibitory inputs primarily from either the ipsilateral or contralateral utricle. These data indicate that bilateral inputs from the utricle represent a large part of excitatory and inhibitory synaptic inputs to vestibulospinal neurons.

**Figure 3:**
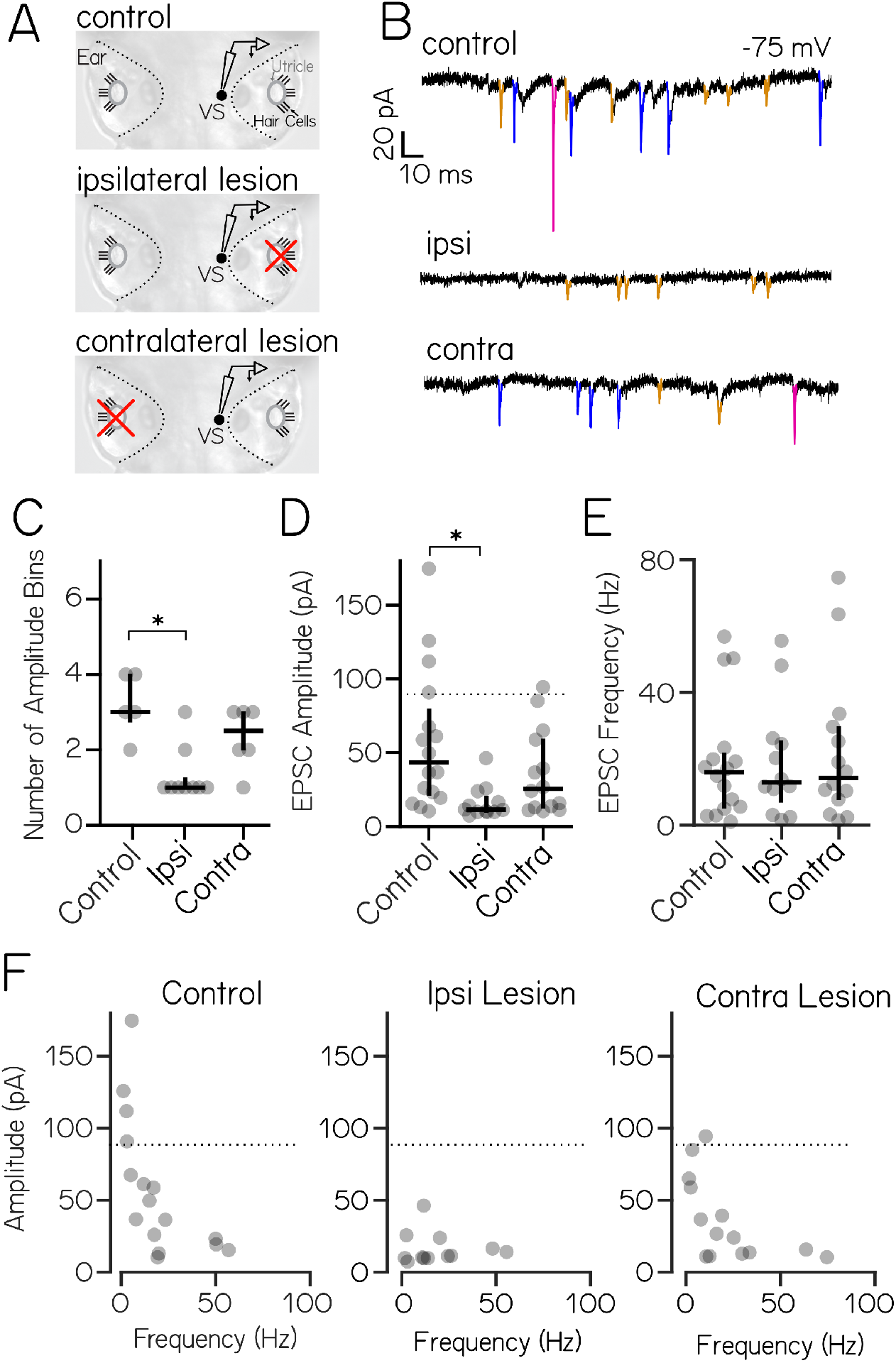
High amplitude spontaneous excitatory inputs originate in the ipsilateral ear. (A) Lesion schematic: The utricle (gray circle) was physically removed (red “x”) either ipsilateral or contralateral to the recorded vestibulospinal neuron (black circle, “VS”) (B) Example current traces from neurons held at −75 mV from control (top), ipsilateral (middle), and contralateral (bottom) experiments; EPSCs in color (C) Number of EPSC amplitude bins per cell (median ± IQR in black) is decreased after ipsilateral, but not contralateral lesion (D) EPSC bin amplitudes (gray circles, median ± IQR in black) are decreased after ipsilateral, but not contralateral, lesion. Dashed line at the top 5% (“high amplitude”) of control EPSCs (E) Frequency of events in EPSC bins (gray circles, median ± IQR in black) is unchanged after ipsilateral or contralateral lesion compared to control cells (F) EPSC amplitude vs. frequency for each bin (gray circles) in control and after ipsilateral/contralateral lesions. High amplitude (top 5% of control) bins are lost after ipsilateral lesion

### Vestibulospinal neurons respond to postural destabilization in a utricle-dependent manner

Our lesion experiments indicate that vestibulospinal neurons are poised to encode posture change through vestibular input from the utricle. We next assayed if vestibulospinal neurons have a sensory-evoked response to postural destabilization, and if so, whether such a response is utricle-dependent. To test this, we manually translated our preparation – in the dark – either along the fore/aft or lateral axis of the fish to provide vestibular stimulation (linear acceleration) to utricular hair cells during whole-cell recordings (Figure 4A). We injected a small amount of depolarizing current so that neurons that were silent at rest would fire at 1Hz to ensure that responses would not be lost due to a high firing threshold. Neurons fired phasically during translation (Figure 4B, top, response to example lateral translation); we quantified their directional tuning by calculating the modulation depth: the strength of the peak phasic response relative to their anti-phase response [50]. We then tested for directional tuning significance in each translational axis (lateral or fore-aft) by comparing a cell’s modulation depth for that stimulus relative to that derived from randomly shuffled data, and categorized each cell as “directional” or “non-directional” for each axis. All control neurons (17/17) responded directionally to lateral stimuli (modulation depth 50.0±28.7 Hz), and most (10/17) were directionally responsive to fore/aft stimuli (modulation depth 23.4±21.7 Hz) (Figure 4C, directional neurons circled). Most neurons were directionally tuned to the peak acceleration of the stimulus towards the contralateral side (6/10) and rostral direction (7/10 neurons) (Figure S3A). EPSC bin amplitudes did not change during translation (paired t-test p=0.6) indicating that the composition of synaptic inputs did not change between rest and stimulation. We conclude that vestibulospinal neuron activity can encode changes in body position.

**Figure 4:**
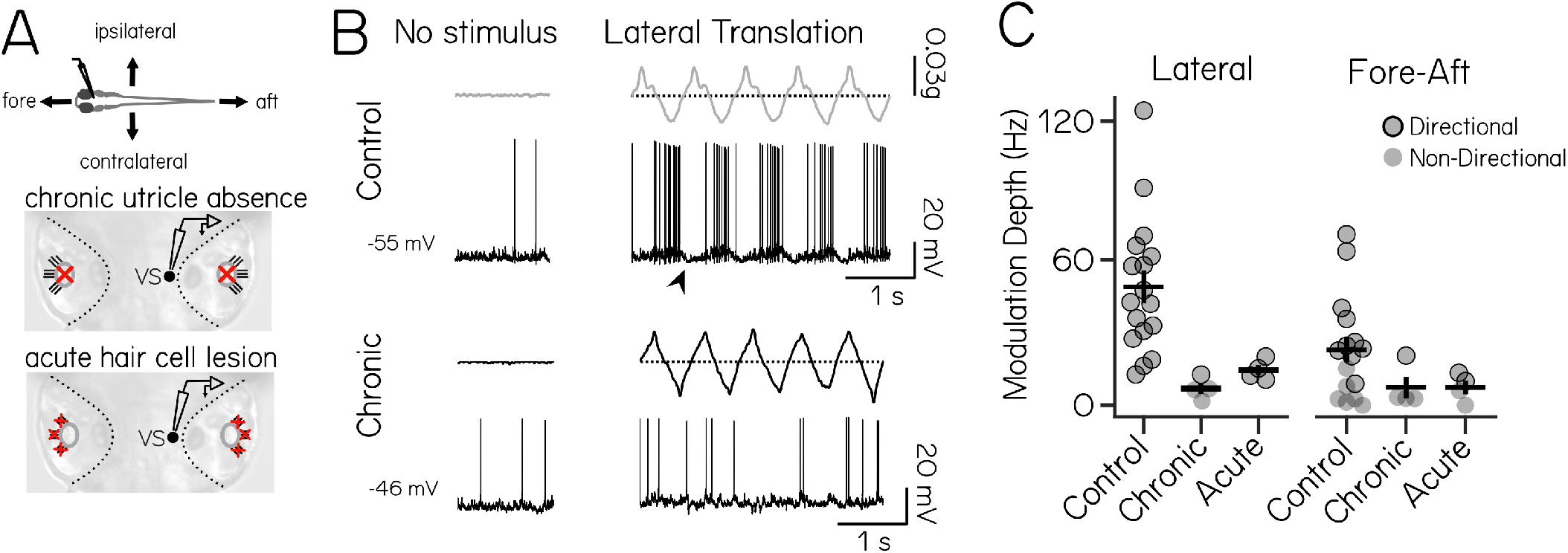
Vestibulospinal neurons encode utricle-derived body translation. (A) Immobilized fish were manually translated in the fore-aft or lateral axes (top). Vestibulospinal neurons were recorded in control and after two manipulations: first, in *otogelin* mutants (middle) that do not develop utricles (red “x”) and second, after chemically-induced hair cell (red “x”) death (bottom) (B) Accelerometer (gray) and voltage trace (black) from a neuron in a control fish (top) showing action potentials and low-amplitude membrane potential fluctuations (arrowhead) in phase with translation. In contrast, activity from a *otogelin* mutant is unaligned with translation (C) Modulation depth of spiking response is disrupted in both the lateral (left) and fore-aft (right) direction after both chronic and acute disruption of the utricle. Gray circles are neurons, black outline denotes statistically significant directional responses

To confirm that action potentials reflected activity originating in the utricular macula, we translated *otogelin* mutants that fail to develop utricles (Fig 4A, chronic utricle absence)[51]. Neurons in mutant fish failed to respond phasically (Figure 4B, bottom). Modulation depth was decreased in mutant fish compared to controls in both the lateral (7.1±4.6 Hz) and fore-aft axis (7.6±8.9 Hz). Among recordings from mutants (n=4) one neuron met our criteria as directionally responsive for lateral translation and a different neuron was directionally responsive for fore/aft translation, but modulation depth was low in both (Figure 4C). Neurons in mutant fish still received normal levels of spontaneous synaptic excitatory input (Figure S4), suggesting that the absence of stimulus response is not due to loss of synaptic input but instead reflects loss of tuned input. These data strongly suggest that the bulk of directionally-sensitive inputs to vestibulospinal neurons originates from the utricle.

To control for compensatory mechanisms in *otogelin* mutants, we measured vestibulospinal neuron responses to translation after acute chemo-ablation of inner-ear hair cells (Figure 4A, acute hair cell lesion; Figure S4A). Modulation depth was also reduced dramatically in both the lateral (14.8±4.3 Hz) and fore-aft axes (7.5±5.9 Hz) after acute hair cell lesion (Figure 4C). Across both acute and chronic (*otogelin*) utricle manipulations, modulation depth was strongly affected by lesion condition, but not stimulus direction (Two-Way ANOVA, main effect of lesion condition F_2,13_=8.5, p=0.0008; main effect of stimulus direction F_1,43_=2.1, p=0.16; interaction effect of lesion condition and stimulus direction F_2,43_=1.6, p=0.22) with lower modulation depth in chronic (Tukey’s posthoc test, p=0.004) and acute conditions (Tukey’s posthoc test, p=0.014) compared to controls. Acute lesions did not decrease the fraction of neurons directionally responsive to lateral and fore-aft stimuli (100% lateral directional, 50% fore-aft directional, n=4), but the strength of tuning was low (lateral = 14.8±4.3 Hz; fore/aft = 12.0±2.5 Hz) (4C, directional neurons circled). Finally, after acute hair cell loss, spontaneous excitatory input onto vestibulospinal neurons was significantly decreased but not completely removed (Figure S4). Taken together with the recordings from *otogelin* mutants, we conclude that utricular input is required for the bulk of the normal phasic response to translation in vestibulospinal neurons.

We observed that the baseline membrane potential of vestibulospinal neurons fluctuated systematically during both fore-aft and lateral translation (Figure 4B, arrowhead). To determine if this low-frequency fluctuation reflected synaptic inputs, we performed voltage-clamp recordings during translation following utricular perturbations. Fluctuation amplitude was reduced or eliminated after utricular manipulations (Figure S3B-C) (Two-Way ANOVA, main effect of lesion condition F_2,43_=8.4, p=0.0008; main effect of stimulus direction F_1,43_=1.04, p=0.31; interaction effect of lesion condition and stimulus direction F_2,43_=0.91, p=0.41) with significantly lower amplitude fluctuation in chronic (Tukey’s posthoc test, p=0.0012) conditions (Tukey’s posthoc test, p=0.08). Our data indicate that membrane potential fluctuations reflect changes to synaptic inputs derived from the utricle. We next examined the contributions of excitatory and inhibitory synaptic inputs to membrane potential fluctuations. EPSC rate was correlated (R^2^=0.36 lateral, 0.45 fore/aft) with membrane potential for both stimulus directions (Figure S3D). Although IPSC rate was modulated by translation (Figure S3E-F) the challenge of recording IPSCs during translation precluded a similar analysis. We nonetheless observed that excitatory and inhibitory inputs to a given neuron could be either cotuned or oppositely-tuned (n=2) (Figure S3E-G). Thus inhibitory inputs could contribute to membrane potential oscillation, and presumably action potential rate, during translation. Taken together, our data link vestibulospinal neuron output (action potentials following translation) to utricle-derived excitatory and inhibitory synaptic input.

### Vestibulospinal neurons mediate specific corrective behaviors that maintain balance

Our anatomical and electrophysiological studies indicated that the larval zebrafish vestibulospinal circuit is poised to regulate posture. To illuminate the specific vestibulospinal contributions to postural stability, we adopted a loss-of-function approach. *Tg(nefma:GAL4);Tg(UAS:EGFP)* reliably labels 30±6 vestibulospinal neurons per brain hemisphere (61% of the total population, Figure S5A), of which we could routinely ablate 23 (±5). Photoablations killed neurons without damaging surrounding cells or neuropil (Figure 5A). Photoablated neurons remained absent from larvae 3 days after the lesion occurred suggesting no recovery or regeneration occurred in these cells (1.7±1.5 cells in lesioned hemisphere vs 14.7±2.4 control hemisphere; paired t-test p=1.7*10^−6^).

**Figure 5:**
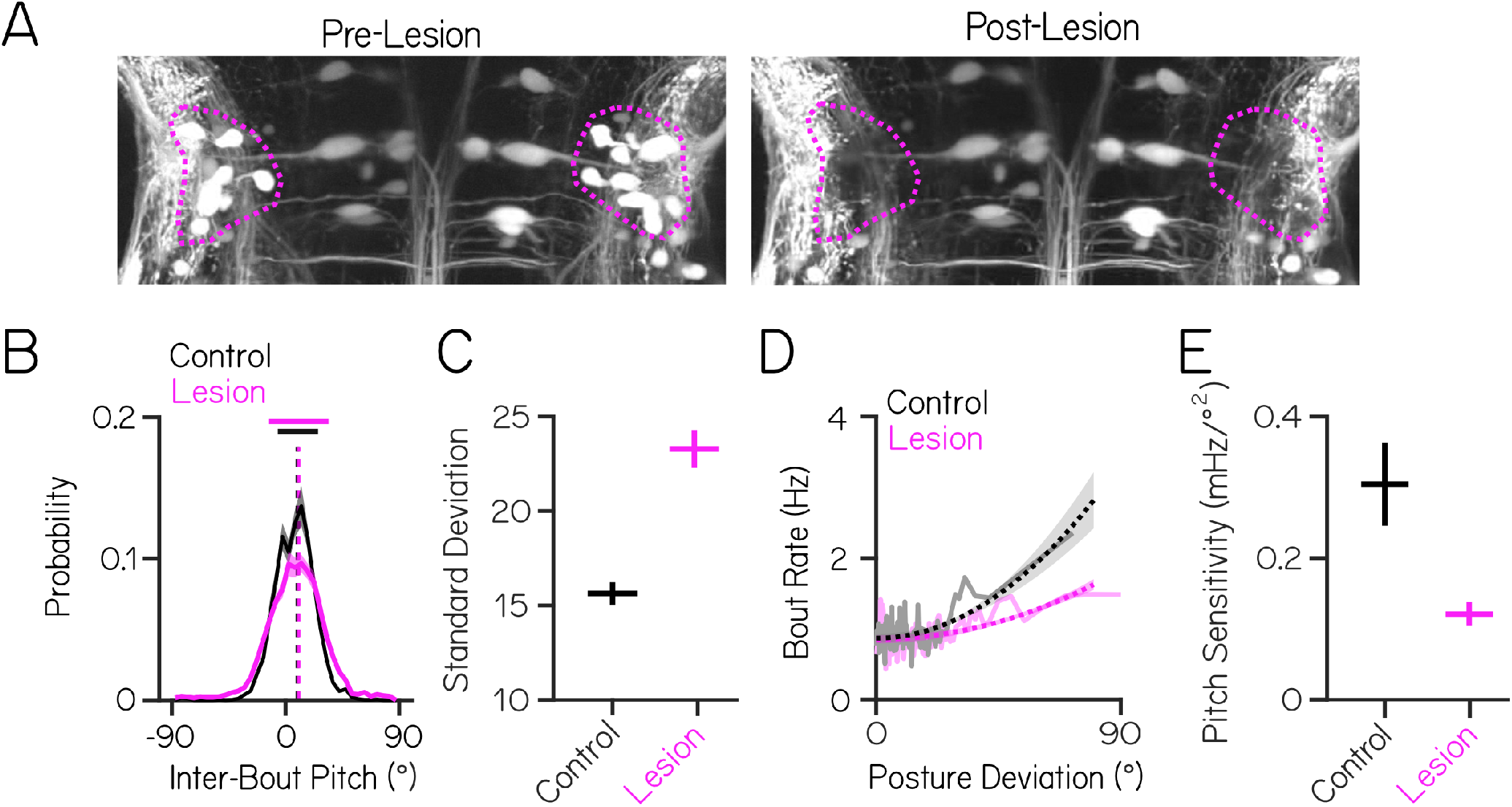
Vestibulospinal neurons contribute to postural stability and movement timing, but not postural set point. (A) Images before and after targeted photoablations of genetically-labeled vestibulospinal neurons show selective loss of fluorescent somata in outlined area (magenta) (B) Distributions of observed pitch show no change to average posture (dashed vertical lines) but greater variability (solid horizontal lines ± 1 SD) between control fish (black) and lesioned siblings (magenta) (C) Postural variability (standard deviation of pitch) is greater in lesioned fish than sibling controls; lines are mean ± jacknifed standard deviation (D) Movement timing (bout rate as a function of deviation from preferred posture) in lesioned fish (magenta) and sibling controls (black). Solid lines are binned raw data, dashed lines are parabolic fits to raw data ± 1 SD (E) Pitch sensitivity (parabolic steepness between posture deviation and bout rate) is lower in lesioned fish than sibling controls)

Using an automated machine vision-based free-swimming assay [37–39], we measured posture in the pitch axis as 97 lesioned fish and 76 control siblings swam in the dark (n = 9 clutch replicates, 10±6 fish per experiment). Lesioned fish exhibited a broader distribution of inter-bout pitches than their control siblings (15.6±0.6 vs 23.3±1.1; paired t-test p=1.6*10^−35^, Figure 5B). Importantly, while both lesioned and control siblings maintained their posture slightly nose-up from horizontal (mean 9.8±0.8° vs. 10.4±0.8°, Figure 5B) lesioned fish are less likely to be found close (±5°) to their preferred posture (23.9% of interbout observations lesion vs 18.3% control) and more likely to be found at eccentric (> ±45°) nose-up and nose-down postures that control fish rarely adopt (5.3% lesion vs 0.7% control). Lesions did not affect basic kinematic properties such as bout speed, bout duration, or bout frequency (Table 2). Consistent with the continued absence of fluorescent neurons after lesions, behavior did not improve across the 48 hour period of the assay (standard deviation of pitch distribution: 23.4±1.2 Day 1 vs 22.9±1.8 Day2, paired t-test p=0.08; pitch sensitivity: 0.14±0.02 Day 1 vs 0.11±0.03 Day 2, paired t-test p=0.005). We conclude that loss of vestibulospinal neurons disrupts regulation of body posture in the pitch axis, but not the preferred posture or swim kinematics.

**Table 2:**
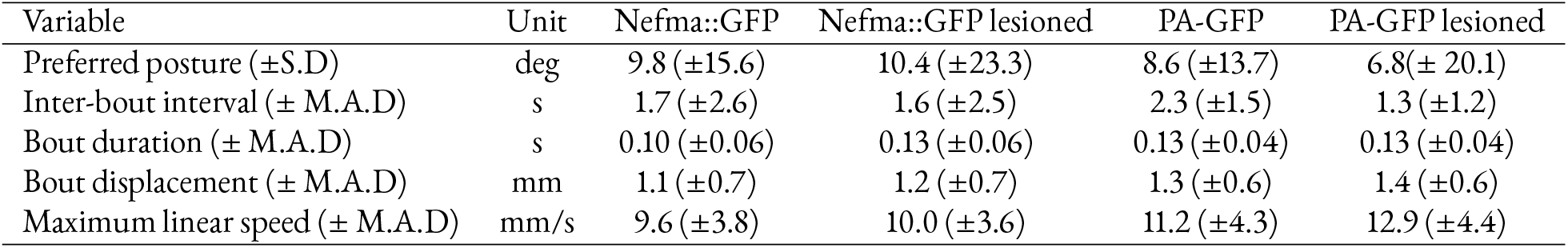
Behavioral properties

| Variable | Unit | Nefma::GFP | Nefma::GFP lesioned | PA-GFP | PA-GFP lesioned |
| --- | --- | --- | --- | --- | --- |
| Preferred posture (±S.D) | deg | 9.8 (±15.6) | 10.4 (±23.3) | 8.6 (±13.7) | 6.8 (± 20.1) |
| Inter-bout interval (± M.A.D) | s | 1.7 (±2.6) | 1.6 (±2.5) | 2.3 (±1.5) | 1.3 (±1.2) |
| Bout duration (± M.A.D) | s | 0.10 (±0.06) | 0.13 (±0.06) | 0.13 (±0.04) | 0.13 (±0.04) |
| Bout displacement (± M.A.D) | mm | 1.1 (±0.7) | 1.2 (±0.7) | 1.3 (±0.6) | 1.4 (±0.6) |
| Maximum linear speed (± M.A.D) | mm/s | 9.6 (±3.8) | 10.0 (±3.6) | 11.2 (±4.3) | 12.9 (±4.4) |

Two key computations regulate postural stability in the pitch axis [37, 38]. Larvae (1) preferentially initiate swim bouts at eccentric pitches and/or high angular velocities and (2) counter-rotate during the bout to return to their preferred posture. To determine if loss of vestibulospinal neurons interferes with these computations, we first assayed the relationship between movement timing (bout rate) and deviation from preferred posture (Figure 5 D). Lesioned fish showed a marked decrease relative to control siblings (0.12±0.02 vs. 0.30±0.06; paired t-test p=2.5*10^−17^; Figure 5E). Similarly, lesioned fish also showed a weaker relationship between bout rate and angular velocity prior to a bout (nose-up: 0.09±0.01 vs 0.13±0.01; paired t-test p=1.1*10^−19^. Nose-down: −0.09±0.01 vs −0.12±0.01; paired t-test p=3.5*10^−19^). Further, bouts were less corrective for pitch instability (pitch correction gain: −0.28±0.01 vs −0.31±0.01; paired t-test p=9.8*10^−23^).

To determine whether the residual ability to control movement timing reflected an incomplete lesion, we repeated our experiments following optical backfill of all spinal-projecting neurons in the *Tg(α-tubulin:C3PA-GFP)* line (Figure S5B). Lesions removed nearly all vestibulospinal neurons (n=42±10 neurons, n=13 fish) and produced comparable effects on the standard deviation of pitch distribution (13.7 controls vs 20.1 lesions) and on pitch sensitivity (0.40 controls vs 0.18 lesions; paired t-test p=0.002) as lesions targeted in the *Tg(nefma:GAL4);Tg(UAS:EGFP)* background (Figure S5C-F). Lesioned fish had comparably impaired corrective counter-rotations (pitch correction gain: −0.25 controls vs −0.17 lesions; paired t-test p=0.005). We conclude that the remaining ability to control movement timing and the magnitude of corrective counter-rotations likely reflects extra-vestibulospinal contributions to posture rather than an incomplete lesion. Taken together, our data argue that loss of vestibulospinal neurons increases variability in the pitch axis by interfering with both movement timing and corrective stabilization.

To determine the postural impact of disruptions to both movement timing and corrective stabilization, we modeled pitch using a generative model of swimming [37] (Figure 6A). In brief, model larvae are subject to passive destabilization that is partially corrected by stochastic swim bouts whose kinematics and timing are drawn from distributions that match empirical observations. Of twelve data-derived parameters (Table 3), four implement the two relevant computations: three that determine the degree to which bout probability depends on either pitch or angular velocity (bout timing) and one determines the degree to which bouts restore posture (bout correction). Both bout timing and bout correction computations are necessary in the model to generate simulated bouts with a preferred horizontal posture and a low-variability pitch distribution (Figure 6B, in agreement with previous findings[37]. Our model could recapitulate the distributions of posture and swim bout timing using parameters derived from control data (Figure 6C-D). In contrast, using parameters derived from lesion data broadened the distribution of pitch angles (Figure 6E) as seen after vestibulospinal lesions. If the vestibulospinal contribution to pitch axis stability reflects disruptions to movement timing and corrective stabilization, then observed changes to both (but not either alone) should explain the increased variability in posture. For each computation, we systematically replaced the relevant parameters derived from control data with those from lesion data. Neither parameter set alone could recapitulate the variability observed in the lesion model (Figure 6E), but together both could. Our model supports the conclusion that the observed disruptions to both movement timing and corrective stabilization together encapsulate the behavioral consequences of vestibulospinal lesions.

**Figure 6:**
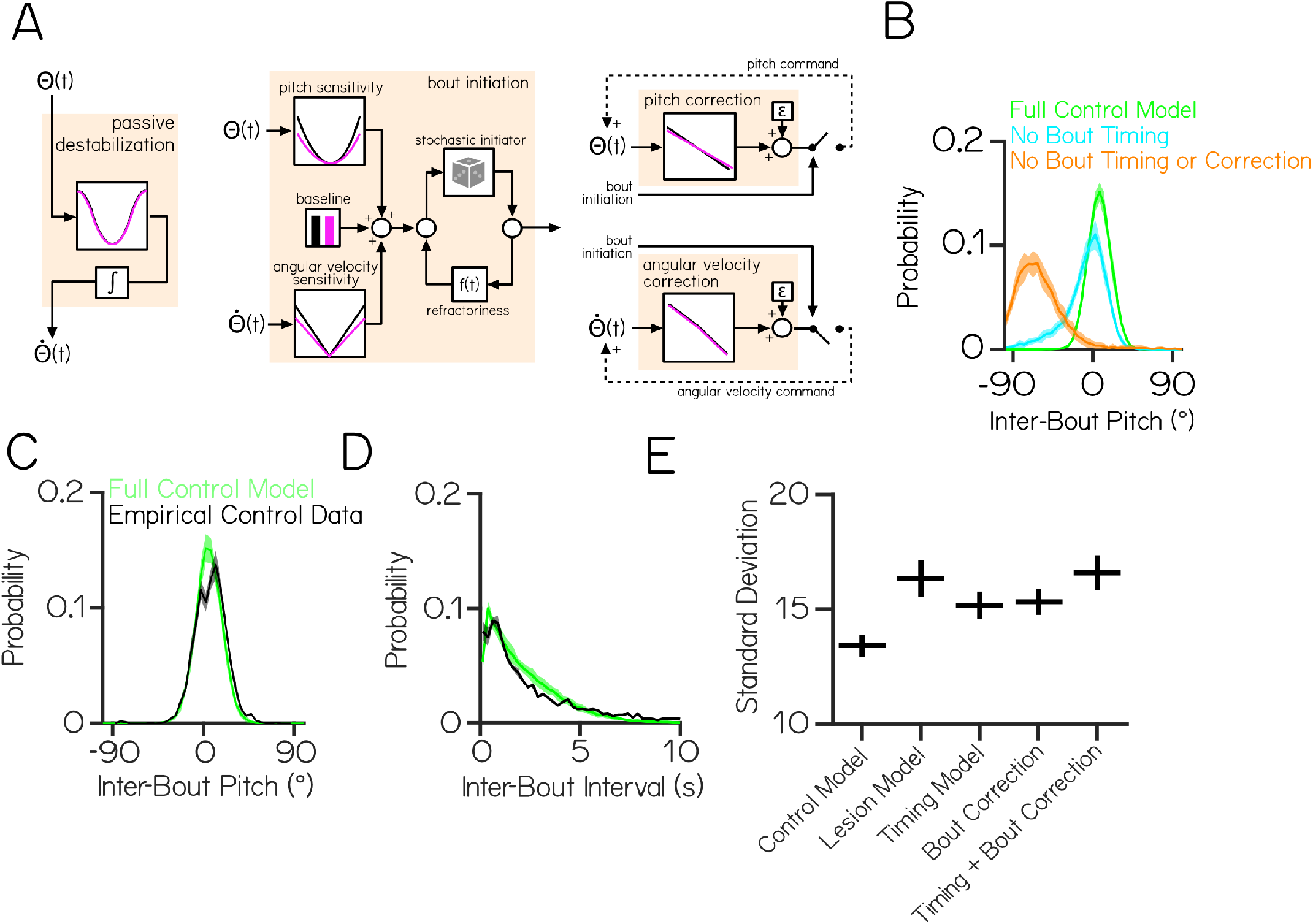
Behavioral modeling shows that increased postural variability following lesions emerges from combined impairments to swim timing and corrective capacity. (A) A generative model of swimming adapted from previously published work [37], consists of four computations (tan boxes) determining swim timing, pitch angle, and angular velocity. Each computation contains one or more condition-specific parameters, shown by plots with a black (control) or magenta (vestibulospinal lesion) line (B) Probability distributions of simulated pitch angles in full control model (green), bout timing null model (cyan), and full null model (orange) (C) Probability distributions of observed pitch angles between swim bouts and (D) inter-bout intervals for empirical control data (black) and simulated control fish (green) (E) Standard deviation of simulated pitch probability distributions ±S.D. using different combinations of condition-specific parameters in the computations shown in (A)

**Table 3:**
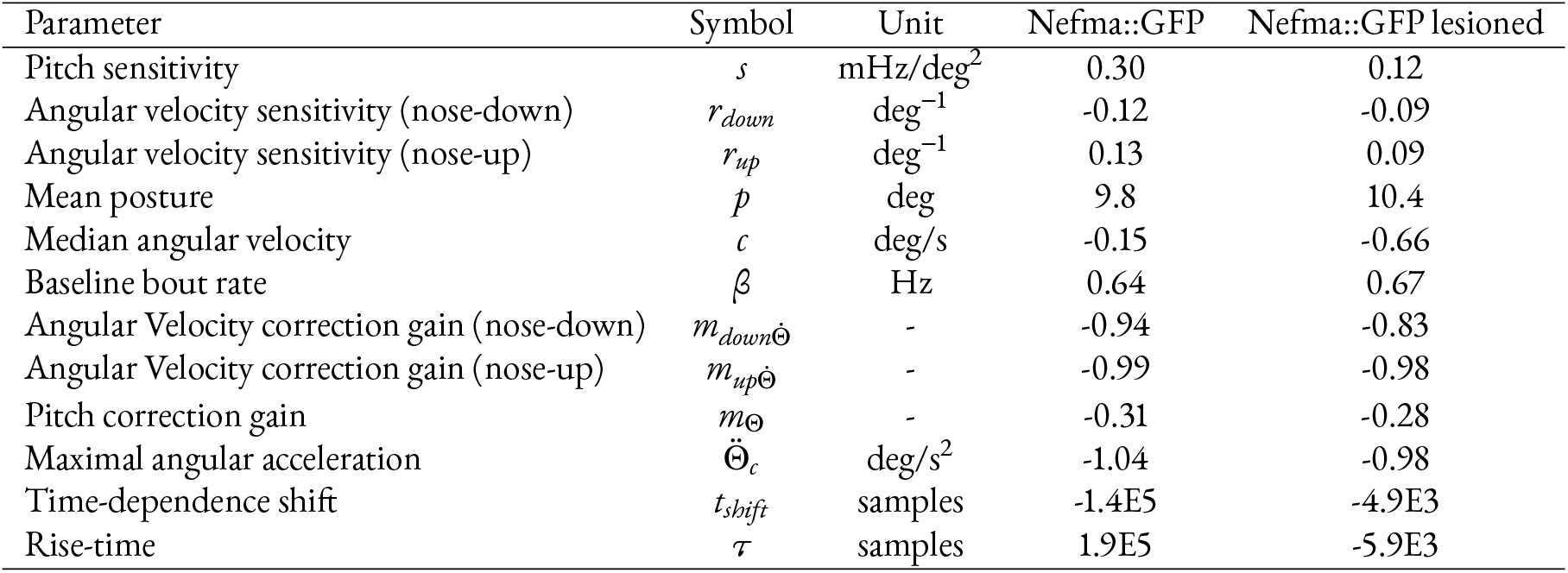
Modeling parameters

### Vestibulospinal neurons mediate fin/body coordination

Zebrafish use vestibular information to coordinate fin and trunk movements to climb in the water column [39]. We therefore hypothesized that vestibulospinal neurons might contribute to this fin-body coordination. Fin use is estimated as the difference between the body angle of the fish and the trajectory of the fish (“attack angle”) (Figure 7A); the slope of the relationship between postural change during a bout and attack angle reflects the degree of fin engagement during climbing. Vestibulospinal lesions decreased this slope (1.2±0.14 vs 1.7±0.14; paired t-test p=7.7*10^−21^) (Figure 7B-C). Our data supports a role for vestibulospinal neurons in coordinating paired appendages (fins) and body (trunk) movements during vertical navigation.

**Figure 7:**
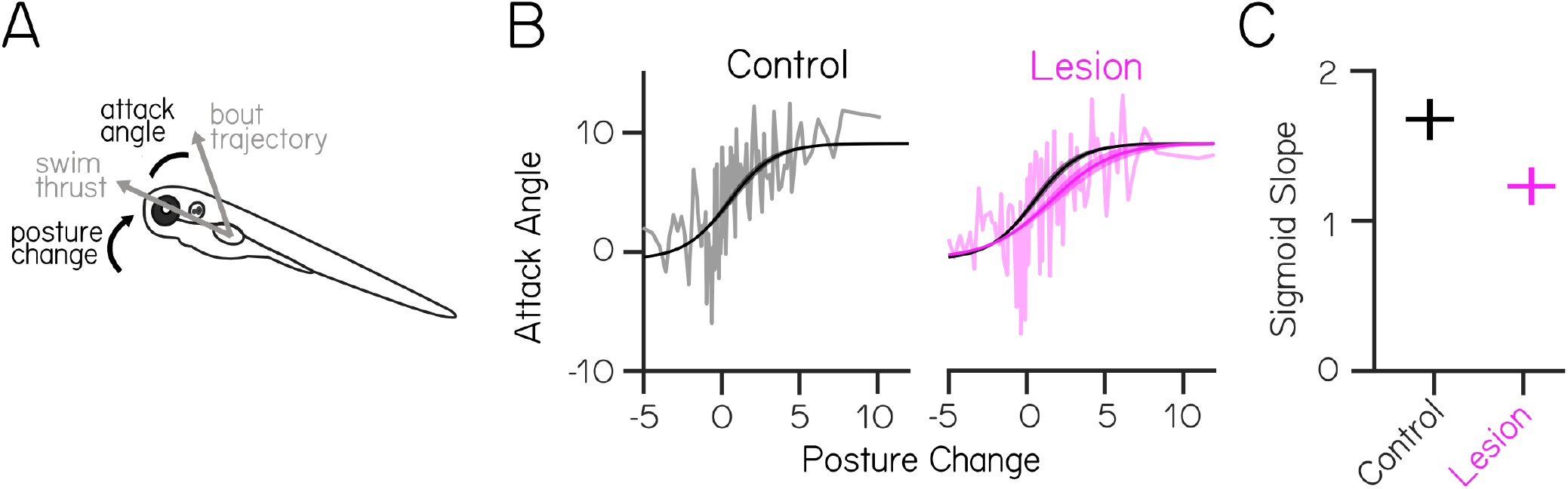
Vestibulospinal neurons are necessary for coordination during vertical climbs. (A) Schematic illustrating fin/body coordination during climbing. Larvae climb by coordinating axial rotation (“posture change”) and forward thrust with fin movements. The fin contribution can be derived from the attack angle, or the difference between the thrust vector and observed bout trajectory (B) Attack angle as a function of posture change in lesioned fish (magenta) and sibling controls (black). Solid lines are binned raw data, dashed lines represent sigmoidal fits ± 1 SD (C) Sigmoid slope decreases in lesioned fish

## Discussion

Vestibulospinal neurons are an evolutionarily ancient population long thought to play a role in balance regulation. Here we use the larval zebrafish as a model to understand the development, synaptic architecture, and behavioral contribution of these neurons. We found that neurons were born and capable of integrating and responding to sensed instability early in development. We identified utricle-derived excitatory, inhibitory, and extravestibular synaptic inputs and showed that neurons use this utricular input to respond to linear accelerations. Our targeted lesions show that acute loss of vestibulospinal neurons leads to postural instability. We determined that this instability reflects both the failure to initiate movements normally and to restore posture appropriately; *in silico* these impairments can explain postural variability. Finally, we discovered that vestibulospinal neurons contribute to proper fin-body coordination, a key component of vertical navigation. Taken together, our data show how vestibulospinal neurons act as a critical and conserved neural locus to transform sensed instability into stabilizing behavior.

Our findings extend recent work by Liu et. al. examining how larval zebrafish vestibulospinal neurons acquire peripheral selectivity [32] by mapping the inhibitory synaptic inputs and defining the behavioral utility for vestibulospinal activity (Figure 8). Intriguingly, our characterization of excitatory inputs disagrees with theirs, suggesting a potential role for modulation of vestibulospinal inputs. We both observed “binned” excitatory inputs and concluded that large amplitude EPSCs likely originate from ipsilateral utricular afferents. However, unlike their recordings, EPSCs in individual bins here rarely passed a refractory period test, and we saw no evidence for electrochemical synapses. The only major difference between the experimental preparations was that our recordings were performed exclusively in the dark while theirs were in ambient light (M. Bagnall, personal communication). We hypothesize that differences might reflect visual or state-dependent modulation of presynaptic inputs to vestibulospinal neurons. Both visual input [52] and behavioral state[53] can profoundly impact vestibular neuron activity, and presynaptic inputs to vestibular neurons are thought to be the site for visually-guided motor learning [54]. In larval zebrafish, such modulation could originate from visually-responsive [55] dopaminergic neurons that when activated drive vestibulospinal neuron activity [56]. Our work thus builds on and expands models of vestibulospinal circuit organization to offer a tractable way to understand – at a synaptic level – how extra-vestibular information influences sensed imbalance.

**Figure 8:**
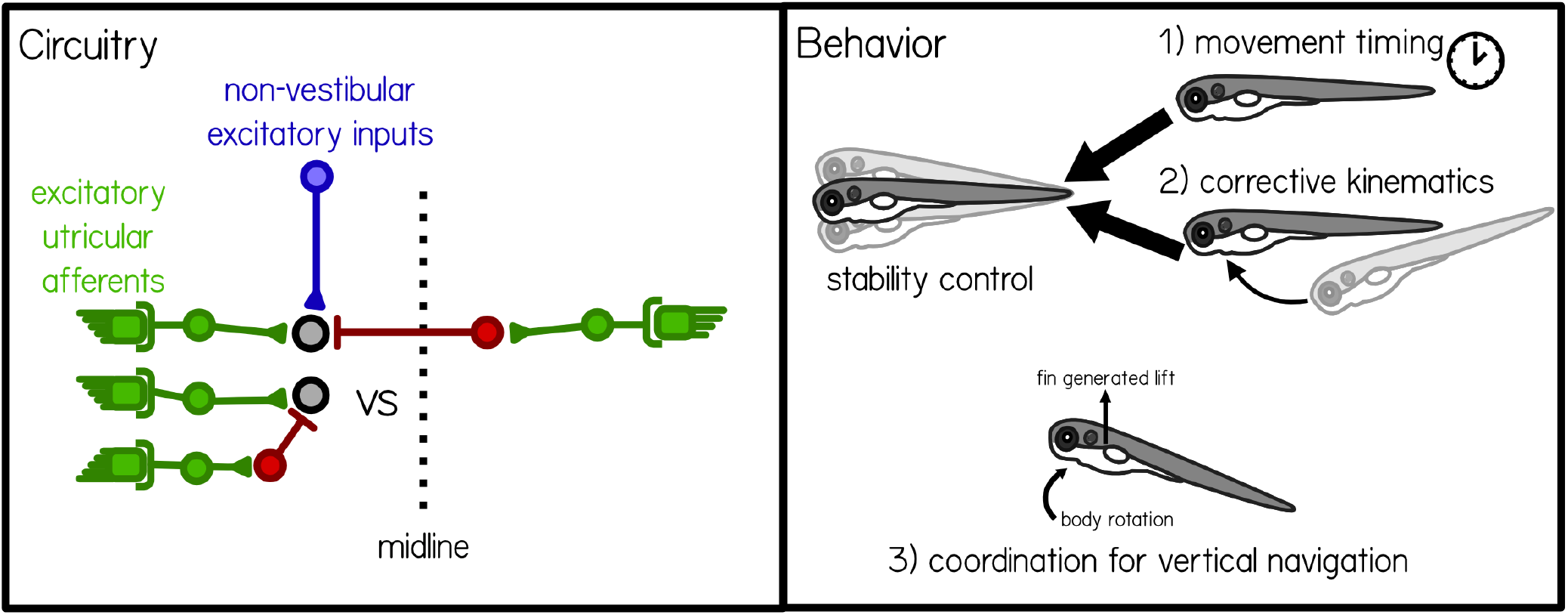
Synaptic architecture and behavioral contributions of vestibulospinal neurons. Circuitry: Vestibulospinal cells receive convergent high amplitude excitatory inputs (green) from irregular afferents originating with the ipsilateral utricle (see also [32]), low-amplitude excitatory inputs (blue) from extra-vestibular sources and inhibitory inputs (red) from either the ipsilateral or contralateral utricle. Behavior: Vestibulospinal cells are involved with three key computations. The first two, movement timing and corrective kinematics together allow fish to maintain postural stability. The third allow fish to coordinate fin and body rotations to climb in the water column.

Our lesion data and model argue that during locomotion vestibulospinal neurons specify both the degree of corrective movements and their timing. Further, they do not appear to determine postural set point or locomotor kinematics even though larvae can modulate both [37, 57]. Similar findings were obtained following partial loss of homologous neurons in the lateral vestibular nucleus, which results in loss of hindlimb extension reflexes following imposed destabilization [27, 28]. Movement initiation (premature stepping) has been observed following stimulation of the lateral vestibular nucleus in cats [58, 59], though see [60]. Previous work was limited to restrained or decerebrate animals; here we advance by measuring unrestrained swimming with detrimental impacts to normal movement timing. Notably, neither our lesions nor comparable mammalian experiments produce gravity-blind animals, suggesting parallel means of postural control. In addition to descending tracts, vestibular information reaches the spinal cord through ascending vestibular neurons in the tangential nucleus [49, 61] that in turn synapse on to spinal-projecting midbrain neurons in the nucleus of the medial longitudinal fasciculus responsible for swim kinematics [57, 62, 63]. Additionally, it is possible that information about body posture might derive from non-vestibular sensory feedback [23, 64]. Taken together, our findings extend complementary loss- and gain-of-function experiments in vertebrates and define part of the neural substrate for turning sensed imbalance into corrective behaviors.

In addition to regulating computations responsible for postural control, we discovered that loss of vestibulospinal neurons disrupts coordinated fin/body movements zebrafish use to navigate vertically in the water column. Considerable evidence indicates that pectoral fins are evolutionary predecessors to tetrapod forelimbs [65] that are driven by molecularly-conserved pools of motor neurons capable of terrestrial-like alternating gait [66]. Might vestibulospinal-mediated coordination of trunk and limbs be similarly conserved? Ancient vertebrates without paired appendages such as lampreys have homologous vestibulospinal neurons, but the projections of these neurons terminate in the most rostral portions of the spinal cord and have been postulated to be important in turning but less crucial for the maintenance of posture [67, 68]. In contrast, vestibular-driven movements of the pectoral fin can be elicited in elasmobranches [69]. Vestibulospinal neurons in frogs form a key part of the circuit that stabilizes posture at rest and coordinates trunk and hindlimb effectors for balance [70]. In terrestrial vertebrates, postural stability relies on coordination of “anti-gravity” extensor muscles in the trunk and limbs [71–74]. Mammalian vestibulospinal neurons innervate spinal regions that control both [7–9, 24, 25, 75, 76]. Our finding that vestibulospinal neurons coordinate fin/trunk movements thus strengthens the proposal that the vestibulospinal circuit serves fundamentally similar roles across disparate body plans and locomotor strategies.

As they grow, larval fish improve their use of vestibulospinal-dependent computations to stabilize posture and navigate [37–39]. However across our population we did not observe concomitant changes to intrinsic or synaptic properties, and the hallmarks of a mature circuit (bilateral excitatory and inhibitory inputs, acceleration-tuned outputs) were present throughout our recordings. We hypothesize that cerebellar regulation of vestibulospinal activity, and not intrinsic maturation of vestibulospinal neurons, underlies behavioral improvements. The cerebellum is well-known to make direct synaptic contact with vestibulospinal neurons in frogs and mammals [12, 42, 77, 78] and is critical for proper adaptation of vestibulospinal reflexes [79–81]. In the larval fish, vestibulocerebellar efferents target vestibulospinal neurons, with marked outgrowth during the period of behavioral improvement [82]; zebrafish Purkinje cells are electrophysiologically mature, responsive, and necessary for motor learning [83, 84] during this time. Vestibular stimulation results in considerable cerebellar activity [33, 34], consistent with its role in coordination of pectoral fin and body posture during navigation in larval fish [39]. The cerebellum is therefore poised to refine vestibulospinal activity to permit balance to mature.

Though vestibulospinal neurons were first described over 150 years ago [85], they remain the focus of active interest today [23, 27, 28, 32, 70]. Here we examined vestibulospinal neurons across many levels, combining new electrophysiological data and precise loss-of-function perturbations with the comparatively simple and well-defined physics of underwater locomotion. Our work makes two specific contributions: First, we delineate a synaptic blueprint from the inner ears to vestibulospinal neurons that encodes sensed imbalance. Second, we show that vestibulospinal neurons contribute to movement timing and corrective kinematics, as well as trunk/paired appendage coordination. These behaviors are not only fundamental for proper posture and locomotion, but each improves with age [37, 39]. Therefore we propose that, in addition to elucidating the synaptic architecture for specific balance behaviors, our findings are foundational for future studies into the neuronal mechanisms underlying vertebrate postural and locomotor development.

## Materials and Methods

### Fish Care

All procedures involving zebrafish larvae (*Danio rerio*) were approved by the Institutional Animal Care and Use Committee of New York University. Fertilized eggs were collected and maintained at 28.5°C on a standard 14/10 hour light/dark cycle. Before 5 dpf, larvae were maintained at densities of 20-50 larvae per petri dish of 10 cm diameter, filled with 25-40 mL E3 with 0.5 ppm methylene blue. After 5 dpf, larvae were maintained at densities under 20 larvae per petri dish and were fed cultured rotifers (Reed Mariculture) daily.

### Fish Lines

Experiments were done on the mitfa-/- background to remove pigment. Larvae for photoconversion experiments were from the *Tg(elavl3:Kaede)^rw0130a^* [86] background. Larvae for vestibulospinal lesions were labeled with *Tg(nefma:GAL4;UAS:GFP)* [32]. Photoconverted larvae used for additional vestibulospinal lesions were from the *Tg(α-tubulin:C3PA-GFP)* [49] background. For chronic bilateral utricular lesions, fish with homozygous recessive loss-of-function mutation of the inner ear-restricted gene, *otogelin* (otog-/-), previously called *rock solo*^AN66^ [51] were visually identified by a lack of utricular otoliths.

### Spinal Photoconversions and Analysis

Kaede- or PA-GFP-positive larvae were raised in a dark incubator to prevent background photoconversion. For photoconversion, larvae were anesthetized with 0.2 mg/mL ethyl-3-aminobenzoic acid ethyl ester (MESAB, Sigma-Aldrich E10521, St. Louis, MO) and mounted laterally in 2% low-melting temperature agarose (Thermo Fisher Scientific 16520). Using a Zeiss LSM800 confocal microscope with 20x objective (Zeiss W Plan-Apochromat 20x/1.0 DIC CG=0.17 M27 75mm), the spinal cord between the mid-point of the swim bladder and the caudal-most tip of the tail was repeatedly scanned with a 405 nm laser until fully converted. For anatomical characterization of vestibulospinal neurons, tail photoconversion of the *Tg(elavl3:Kaede)* transgenic fish was performed 4-24 hours prior to imaging at 5 dpf. For retrograde labelling of vestibulospinal neurons used for photoablations, the spinal cords of *Tg(α-tubulin:C3PA-GFP)* larvae were converted using the same protocol at 6 dpf. To allow the converted fluorophore to diffuse into cell bodies, all fish were removed from agarose after photoconversion and raised in E3 in a dark incubator.

Anatomical image stacks of photoconverted Kaede-positive vestibulospinal neurons in the hindbrain were taken at 5 dpf using a Zeiss LSM800 confocal microscope with 20x objective. Spinal-projecting cells in rhombomere 4 were defined as vestibulospinal neurons if they were located mediolaterally between the Mauthner cell body and the otic vesicle, and if they were located dorsoventrally between ± 30 *μ*m from the tip of the Mauthner lateral dendrite. To generate a spatial map of vestibulospinal cell position across fish, image stacks were first processed using Fiji [87] to split stacks into 6 *μ*m sub-stacks dorsoventrally for each hemisphere of the hindbrain.

Vestibulospinal cell position was then registered manually in Adobe Illustrator by aligning image stacks across fish to a reference stack. Images were aligned across fish using the Mauthner cell body as an anatomical guide, such the lateral-most point of the Mauthner dendrite was defined as 0 *μ*m dorsoventrally. Vestibulospinal cell locations were marked for each fish in reference to the Mauthner lateral dendrite for both the right and left hemispheres. Dorsoventral position of vestibulospinal cells was defined as the 6 *μ*m sub-stack in which the cell was most strongly in-focus.

### Spinal Backfills

To label spinal-projecting neurons in the hindbrain, larvae were anesthethized in 0.2 mg/mL MESAB and mounted laterally in 2% low-melting temperature agarose. Agarose was removed above the spinal cord at the level of the cloaca. An electrochemically sharpened tungsten needle (10130-05, Fine Science Tools, Foster City, CA) was used to create an incision in the skin, muscles, and spinal cord of the larvae. Excess water was removed from the incision site, and crystallized dye (dextan-conjugated Alexa Fluor 647 dye (10,000 MW, ThermoFisher Scientific D-22914) or dextan-conjugated Alexa Fluor 546 dye (10,000 MW, ThermoFisher Scientific D-22911)) was applied to the incision site using a tungsten needle. Larvae were left in agarose for at least 5 minutes after dye application before E3 was applied and the fish was removed from agarose. Fish were allowed to recover in E3 for 4-24 hours before imaging.

### Kaede Birthdating and Analysis

Kaede-positive larvae were raised in a dark incubator to prevent background photoconversion. At 18, 20, 24, 28, or 32 hours post-fertilization, larval fish were exposed to a 405nm LED for 3-5 minutes in a custom-built apparatus to completely photoconvert Kaede from green to red ubiquitously. Larvae were returned to a dark incubator and raised until 4 dpf, when vestibulospinal neurons were then labeled with a spinal backfill as above. Fish recovered for 24 hours and were imaged using a Zeiss LSM800 confocal microscope with 20x objective.

Confocal images were analyzed across three channels (converted Kaede, unconverted Kaede, backfill dye) by first identifying vestibulospinal neurons in the backfill channel, and subsequently determining if the cell expressed converted Kaede. Converted Kaede was taken as evidence that the vestibulospinal cell had been born at the time of photoconversion. To differentiate spontaneous from induced photoconversion, reference images were taken from fish that had not been exposed to UV light. A threshold pixel intensity was defined as that which removed all visible expression in the red Kaede channel for each time point. The threshold was then used to define a minimum red pixel saturation level in photoconverted fish to differentiate real Kaede conversion from background conversion. For each brain hemisphere of each fish, total number of labeled vestibulospinal cells was counted, as well as the number of converted and unconverted vestibulospinal cells. Percent of cells converted was calculated for each brain hemisphere where more than 4 vestibulospinal cells were labeled by the backfill. Mean percent was calculated across brain hemispheres for each conversion time point.

### Electrophysiology

Larval zebrafish between 3-15 dpf were paralyzed with pancuronium bromide (0.6mg/mL) in external solution (in mM: 134 NaCl, 2.9 KCl, 1.2 MgCl_2_, 10 HEPES, 10 glucose, 2.1 CaCl_2_) until movement ceased. Fish were then mounted dorsal-up in 2% low-melting temperature agarose and a small incision was made in the skin above the cerebellum. Pipettes (7-9 MOhm impedance) were lowered to the plane of the Mauthner cell body, illuminated by infrared (900nm) DIC optics. Putative vestibulospinal neurons were targeted for patching using soma size and proximity to the Mauthner lateral dendrite as guides. Initial targeting conditions were determined with reference to cells labelled by spinal backfills. Subsequent recordings were determined to be from vestibulospinal neurons by post-recording analysis of anatomical morphology, confirming a single ipsilateral descending axon using either widefield fluorescence or confocal microscopy.

All electrophysiological measurements were made in the dark. Pipettes were filled with dye (Alexa 647 hydrazide, Thermo Fisher A20502) in the internal solution (in mM: 125 K-gluconate, 3 MgCl_2_,10 HEPES, 1EGTA, 4 Na_2_-ATP). For recordings with voltage clamp trials at 0 mV holding potential, pipettes were filled with an internal solution to prevent action potentials (in mM: 122 CsMeSO3, 5 QX-314 Cl, 1 TEA-Cl, 3 MgCl_2_,10 HEPES, 1 EGTA, 4 Na_2_-ATP). During trials to determine rheobase and maximum firing frequency, cells were injected with 3 pulses (0.5 sec pulse duration) of decreasingly hyperpolarizing current, and 7 pulses (0.5 sec pulse duration) of increasingly depolarizing current. The magnitude of current steps were scaled for each cell until spikes were seen during the strongest depolarizing current. The average current step size for control vestibulospinal cells was 21.75 pA (range 12-46 pA) with average peak depolarizing current of 152.25 pA (range 84-322 pA). In current clamp trials during linear acceleration stimuli, cells were often injected with an offset depolarizing current (range 0-292.7 pA, mean 46.8 pA across conditions; 57.6 pA control, 32.8 pA acute bilateral utricle removal, 23.8 pA chronic bilateral utricle removal) until spontaneous action potentials were seen during the baseline period without translation.

### Linear Translation Stimulus

Fish were mounted dorsally on an air table (Technical Manufacturing Corporation) with handles (McMaster Carr 55025A51) mounted underneath for manual translation. To produce table oscillations, the air table was manually pushed in either the lateral or forward direction to produce persistent table oscillations. Acceleration traces of table oscillations were measured using a 3-axis accelerometer (Sparkfun, ADXL335) mounted to the air table. Table oscillations persisted for an average of 11.69±5.05 seconds, with a mean frequency of 1.54±0.17 Hz and peak acceleration of 0.91±0.21 g across all trials (n=122 trials (73 lateral; 49 fore-aft)). Lateral translation trials were longer in duration than fore-aft trials (13.02 s lateral vs. 9.71s fore-aft; p=2.98*10^−4^ unpaired t-test), and higher in peak acceleration (1.01g lateral vs 0.75g fore-aft; p=3.02*10^−14^), but did not significantly differ in stimulus frequency from fore-aft stimuli (1.52 Hz lateral vs 1.58 Hz fore-aft; p=0.051).

### Electrophysiology Analysis

Data analysis and modeling were performed using Matlab (MathWorks, Natick, MA, USA). For current clamp recordings, action potentials were identified with reference to a user-defined threshold for each trial. Action potential amplitude was calculated as the difference between peak membrane voltage and a baseline voltage (1 ms prior to event peak). Rheobase for vestibulospinal action potential generation was calculated by fitting a line to firing responses as a function of injected current, limited to all current steps above the minimum current injection which elicited spikes. The linear fit was used to solve for the estimated current necessary for each vestibulospinal cell to fire at 1 Hz.

For voltage clamp recordings, excitatory (EPSC) and inhibitory (IPSC) events were identified through hand-selection of events with a waveform consisting of an initial sharp amplitude rise and exponential-like decay. Exact event times were further identified by local minima or maxima search around hand-selected event times. EPSC amplitudes were determined by subtracting the minima of the event waveform from a pre-event baseline current (0.4 ms before event minima). For EPSCs, amplitude “bins” were assigned manually using the probability distributions of EPSC amplitudes across all trials from the same cell. EPSCs with amplitudes less than 5 pA were excluded from analyses. IPSC amplitudes were determined by taking the difference between the maxima of the event waveform and a pre-event baseline current (2 ms before event maxima).

#### Assessment if EPSCs had a unitary origin

To determine if EPSC bins were derived from a single afferent origin, we calculated the number of within-bin EPSCs that occurred within a 1 ms refractory period. We excluded events that occurred within 0.3 ms of each other from this analysis as manual inspection found that these were usually double-selections of the same synaptic event, rather than two separable events. We only rarely observed vestibulospinal EPSC amplitude bins with zero within-bin violations. To minimize Type II errors from overly strict refractory period violation criteria, we modeled an upper limit on the number of within-bin refractory period violations that we would expect to see from the overlap in EPSC amplitude distributions: The probability distribution of EPSC amplitudes (*I*) of each cell was estimated as as a sum of Gaussian distributions, where the number of distributions was set to the number of EPSC amplitude bins in that cell. Each amplitude bin was fit with 3 free parameters for height (*h*), center (*c*) in pA, and standard deviation (*σ*):

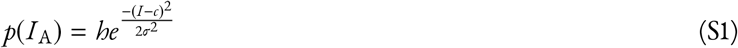

Bin centers (*c*) were constrained to fall within bin amplitude cut-offs. For cells with only one amplitude bin or for the highest amplitude bin, *c* was constrained at the upper bound to be one standard deviation above the peak amplitude probability. Bin heights (*h*) were constrained to be at least half of the maximum value of the empirical probability distribution within the amplitude bin limits. For a given bin of interest (*A*), we used the modeled probability distributions to calculate the number of expected false-positive refractory period violations (an across-bin event pair being falsely counted as a within-bin event pair, *ϕ*_A_) according to the formula:

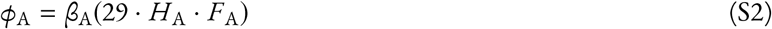

where *β*_A_ is the number of observed EPSC events falling with amplitude bin, *A* (defined by EPSC amplitude thresholds from *a* to *b* pA). The hit event rate (events assigned to bin *A* that truly derived from bin *A*) was calculated by *H*_A_:

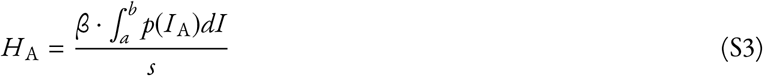

where *β* is the total number of observed EPSCs events across all amplitude bins, and *s* is the trial length in samples. The false-positive event rate for bin *A* (events falsely assigned to bin *A* when they derived from the overlapping tails of Gaussian distributions from other bins) was calculated by *F*_A_:

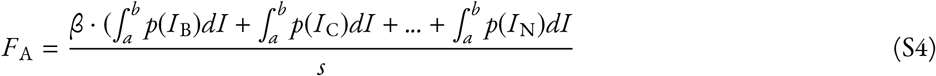

where *p*(*I*_B_) is the estimated probability distribution of the second EPSC amplitude bin, and *p*(*I*_N_) is the n^th^ EPSC amplitude bin.

For each amplitude bin in the cell, *ϕ* was calculated, and compared to the number of observed within-bin violations. To determine whether bins with few to no observed refractory period violations occurred solely due to low event frequency, we calculated the number of expected refractory period violations in frequency-matched randomly generated event trains. EPSC bins were classified as unitary afferent bins if the number of empirical within-bin refractory period violations was fewer than the number of expected violations from bin overlap (*ϕ*) and if the frequency-matched generated controls had at least 1 observed violation.

#### Quantification of sensory responses

Instantaneous spike rates were estimated by computing a peri-stimulus time histogram (PSTH) with a bin width of 1/16 of the cycle length. A cell’s response was defined as the average PSTH across all stimulus cycles in a particular direction (lateral or fore-aft). Modulation depth was determined from this response, defined as the difference between the peak rate and the rate 180° out-of-phase (8 bins away). Cells were considered “directional” if their sensitivity was greater than three standard deviations from the mean modulation depth derived from 100 randomly generated frequency-matched spike trains.

During vestibular stimulation, many cells exhibited low-frequency membrane potential modulations that matched table oscillation frequency. The amplitude of these potential modulations was quantified by first applying a boxcar smoothing filter (width 100ms), then fitting smoothed potential traces during stimulus trials with a sinusoidal function, with 1 fixed parameter for sinusoid frequency set to the stimulus translation frequency, and two free parameters for sinusoid amplitude and DC offset. Occasionally cells that did not display low-frequency membrane potential modulations would have non-zero amplitudes when fit with sinusoids due to membrane potential noise during the stimulation period. To correct for baseline noise, sinusoid amplitude for each cell was calculated by subtracting off the amplitude of a sinusoid fit to membrane potential fluctuations during a baseline period of the recording.

### Utricular Lesions

Chronic bilateral utricular lesions were achieved using mutant larvae (*otogelin*) that do not express *otogelin* and do not develop utricles until 11-12dpf. Acute ipsilateral and contralateral lesions were performed using forceps to rupture the otic capsule and remove the utricular otolith from one ear. This physically removes the sensory organ itself, and likely damages the closely apposed hair cells in the utricular maculae whose spontaneous activity influences vestibular afferents. Acute bilateral lesions were performed through microinjection of 1 mM CuSO4 into both otic capsules to kill hair cells [88], with co-injection of 40 uM FM1-43 dye (Invitrogen T3163) to label hair cell membranes for visualization. After ipsilateral utricle removal and bilateral copper injection, we saw a marked decrease in spontaneous inputs onto vestibulospinal neurons (Figure 3, S4), supporting the hypothesis that these lesions work to impair the firing rate of utricular vestibular afferents. Acute lesions may also impair inner-ear function by diluting the potassium-rich ionic composition of ear endolymph that is critical for hair cell function[89].

### Vestibulospinal Photoablations

Vestibulospinal photoablations were performed on *Tg(nefma:GAL4);Tg(UAS:GFP)* larvae at 6 or 7 dpf for use in behavioral experiments from 7-9 dpf. Additional experiments were performed on *Tg(α-tubulin:C3PA-GFP)* larvae after spinal photoconversion to target a larger number of vestibulospinal neurons. Larvae were anesthetized in 0.2 mg/ml MESAB and then mounted in 2% low-melting point agarose. Photoablations were performed on an upright microscope (ThorLabs) using a 80MHz Ti:Sapphire oscillator-based laser at 920 nm for cell visualization (SpectraPhysics MaiTai HP) and a second, high-power pulsed infrared laser for two-photon mediated photoablation (SpectraPhysics Spirit 8W) at 1040 nm (200 kHz repetition rate, 500 pulse picker, 400 fs pulse duration, 4 pulses per cell over 10 ms) at 25-75 nJ per pulse, depending on tissue depth. Sibling controls were anesthethized for matched durations as lesioned fish. Lesioned and control sibling larvae were allowed to recover for 4-24 hours post-procedure and were confirmed to be swimming spontaneously and responsive to acoustic stimuli before behavioral measurements. Post-lesion survival was high, with only 6/103 fish being excluded from behavioral trials due to failed recovery.

### Behavioral measurement

All behavioral experiments were performed beginning at 7 dpf. *Tg(nefma:GAL4);Tg(UAS:GFP)* experiments were performed on 97 vestibulospinal lesioned larvae, and 76 unlesioned sibling controls (9 paired clutch replicates). *Tg(α-tubulin:C3PA-GFP)* experiments were performed on 17 vestibulospinal lesioned larvae, and 17 unlesioned sibling controls (5 paired clutch replicates). Larvae were filmed in groups of 1-8 siblings in a glass tank (93/G/10 55×55×10 mm, Starna Cells, Inc., Atascadero, CA, USA) filled with 24-26 mL E3 and recorded for 48 hours, with E3 refilled after 24 hours. Experiments were performed in constant darkness.

As described previously[37, 39], video was captured using digital CMOS cameras (Blackfly PGE-23S6M, FLIR Systems, Goleta CA) equipped with close-focusing, manual zoom lenses (18-108 mm Macro Zoom 7000 Lens, Navitar, Inc., Rochester, NY, USA) with f-stop set to 16 to maximize depth of focus. The field-of-view, approximately 2×2 cm, was aligned concentrically with the tank face. A 5W 940nm infrared LED back-light (eBay) was transmitted through an aspheric condenser lens with a diffuser (ACL5040-DG15-B, ThorLabs, NJ). An infrared filter (43-953, Edmund Optics, NJ) was placed in the light path before the imaging lens. Digital video was recorded at 40 Hz with an exposure time of 1 ms. Kinematic data was extracted in real time using the NI-IMAQ vision acquisition environment of LabVIEW (National Instruments Corporation, Austin, TX, USA). Background images were subtracted from live video, intensity thresholding and particle detection were applied, and age-specific exclusion criteria for particle maximum Feret diameter (the greatest distance between two parallel planes restricting the particle) were used to identify larvae in each image. In each frame, the position of the visual center of mass and posture (body orientation in the pitch, or nose-up/down, axis) were collected. Posture was defined as the orientation, relative to horizontal, of the line passing through the visual centroid that minimizes the visual moment of inertia. A larva with posture zero at any given time has its longitudinal axis horizontal, while +90° is nose-up vertical, and −90° is nose-down vertical.

### Behavioral Analysis

Data analysis and modeling were performed using Matlab (MathWorks, Natick, MA, USA). As previously described [37, 39], epochs of consecutively saved frames lasting at least 2.5 sec were incorporated in subsequent analyses if they contained only one larva. Instantaneous differences of body particle centroid position were used to calculate speed. Inter-event intervals (IEIs) were calculated from bout onset times (speed positively crossing 5 mm/sec) in epochs containing multiple bouts, and consecutively detected bouts faster than 13.3 Hz were merged into single bouts.

Numerous properties of swim bouts were calculated. The maximum speed of a bout was determined from the largest displacement across two frames during the bout. Trajectory was calculated as the direction of instantaneous movement across those two frames. Bouts with backwards trajectories (>90° or <-90°) were excluded from analysis. Displacement across each pair of frames at speeds above 5 mm/sec was summed to find net bout displacement. Bout duration was calculated by linearly interpolating times crossing 5 mm/s on the rising and falling phases of each bout. Instantaneous bout rate was calculated as the inverse of the IEI duration. Attack angle was defined as the difference between trajectory and posture of a larva at the peak speed of a bout, such that a larvae pointed horizontally and moving vertically upwards had an attack angle of 90°. Posture change during a bout was defined as the difference in body orientation observed between 25 and 75 ms before peak speed. Pitch angle distributions were computed using inter-bout pitch, or the mean pitch angle across the duration of an IEI. Pitch probability distributions were calculated using a bin width of 5° (ranging from ±90°).

A parabolic function was used to fit the relationship between instantaneous bout rate (*y*) in Hz and deviation from preferred posture(*x*) in degrees, based on the formula

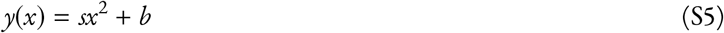

in which *s* gives parabola steepness (pitch sensitivity, in Hz/deg^2^) and *b* gives basal bout rate (Hz). Deviation from preferred posture was itself a function of inter-bout pitch, following the formula:

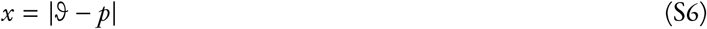

where *ϑ* is the inter-bout pitch (deg) and *p* is the mean inter-bout pitch across all IEIs for each condition. Parameter fits were estimated in Matlab using nonlinear regression-based solver (least-squares estimation). Initial parameter values were *s*=0.001 and *b*=1.

As described previously [39], a logistic function was used to fit the relationship between attack angle (*γ*) and posture change(*r*), based on the formula

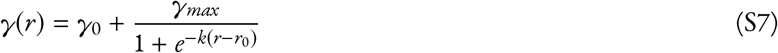

where *γ*_0_ gives the most negative attack angle (in deg), (*γ_max_* + *γ*_0_) gives that largest positive attack angle (in deg), and *k* is the steepness parameter (in deg^−1^). From the derivative of S7, sigmoid maximal slope (found at *r*=*r*_0_) is given by *kγ_max_*/4. Sigmoid center position (*r*_0_) was itself defined from a parameter for rise position (*r_rise_*, posture change at which the sigmoid rises to 1/8 of its upper asymptote:

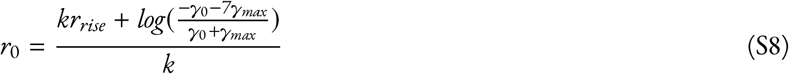

Parameter fits were estimated in Matlab using a nonlinear regression based solver (trust-region-reflective). Initial parameter values were *k*=1 deg^−1^, *γ*_0_=-0.2°, *γ_max_*=10°, and *r_rise_*=-1°. As previously [39], only data from swims that occurred during circadian day (as determined by incubator light cycle) were included. To improve fit, we excluded attack angles > | ± 50°| (1.3% and 1.6% of observations in lesions and controls respectively). To compare sigmoid slope between conditions, *γ*_0_, *γ*_max_, and *r*_rise_ were fixed at means between lesion and control conditions to fit a one-parameter sigmoid.

Due to low bout numbers per experiment (443 ± 271 bouts per clutch for *Tg(nefma:GAL4);Tg(UAS:GFP)* experiments), a single experiment often did not contain enough data to sufficiently sample the entire distribution of postures. Therefore data for all calculations were pooled across all bouts in a given group (lesion or control). Estimates of variation across clutches was performed using jackknifed resampling, where each sample left out two experimental clutch replicates. Paired statistics for behavioral measurements used paired jackknifed sub-samples.

### Behavioral Modeling

We generated condition-specific swimming simulations using a generative swim model described previously [37], with updates to the model to improve the fit to the empirical control dataset collected here. In each condition, 20 simulated fish swam for 3000 seconds with discrete time steps equivalent to those in the captured data (Δ*t* = 25 ms). At each time step (*t*), the pitch (**Θ**(*t*)) is updated due to passive posture destabilization from angular acceleration, with **Θ**(*t*) initialized at a randomly drawn integer from a uniform distribution between ±90°. Angular velocity 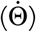 was initialized at 0, and was calculated as the sum of 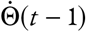 and the integral of angular acceleration, 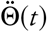:

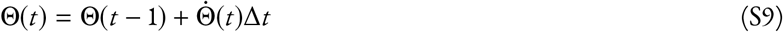

Between swim bouts, simulated larvae were destabilized according to angular acceleration. Angular acceleration varied as a function of pitch in the preceding time step (**Θ**(*t* − 1):

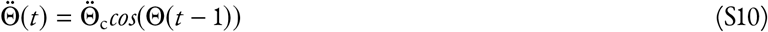

where 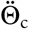 is the maximal empirical angular acceleration observed between bouts for fish each condition, calculated by taking the median of the second differential of smoothed pitch angles between bouts using a moving average filter with a span of 75 ms.

Simulated larval pitch was updated across time according to passive destabilization until a swim bout was initiated. When a bout was initiated, pitch and angular velocity were updated according to condition-specific correction parameters based on empirical swim bout kinematics. Angular velocity correction from swim bouts was corrected by making net angular acceleration across swim bouts 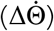 correlated with pre-bout angular velocity 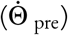. Condition-specific correlations were determined by best-fit lines to empirical data, defined by a slope 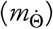 and intercept 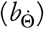 for both positive (nose-up) and negative (nose-down) angular velocities. To reproduce the empirical variability of bout kinematics, net angular acceleration incorporated a noise term 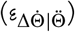 drawn randomly from a Gaussian distribution with mean of 0, and standard deviation 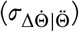 calculated from the empirical standard deviation of 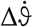 and reduced proportionally by the unexplained variability from the correlation between 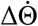 and 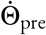.

A bout initiated at time *t* corrected 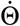 after completion of the bout, 100 ms later (4 time samples, matched to empirical bout duration):

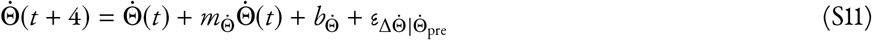

The same approach was used to condition net bout rotation (**ΔΘ**) on pre-bout pitch (**Θ**_pre_) based on a single best-fit line, with a single slope (*m***_Θ_** and intercept (*b***_Θ_**). The corresponding noise term (*ε***_Θ_**) was not reduced by the unexplained variability to better match empirical pitch distributions. For all conditions, the fit line was constrained to the linear portion of the relationship (−30 to +40° pre-bout pitch). If the simulated fish’s posture prior to a bout was outside of this window (only 1% of simulated interbouts), net bout rotation was calculated using *m*_−30_, *b*_−30_ if **Θ** < −30, or *m*_40_, *b*_40_ if **Θ** > 40. To further impose a ceiling on the counter-rotation values attainable by our simulated larvae, a maximum net bout rotation was imposed (**ΔΘ**_max_=±26.3°) based on empirical values; if **ΔΘ** > **ΔΘ**_max_, then **ΔΘ** was set to **ΔΘ**_max_.

Bouts occurred in the model based an internal state variable representing the probability of bout initiation (*P*_bout_). Bout initiation was calculated as the sum of a non-posture dependent baseline bout rate variable (*β*), a **Θ**-dependent bout rate, and 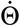-dependent bout rate:

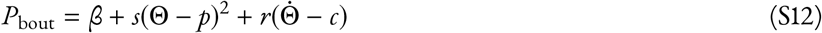

Bout rate as a function of pitch was calculated by fitting a parabola to the relationship between instantaneous bout rate and deviation from mean posture (**Θ** − *p*) to get a pitch sensitivity parameter (*s*) for each condition. Bout rate as a function of angular velocity was calculated by fitting a line to the correlation between instantaneous bout rate and mean-subtracted angular velocity 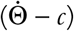 to get a slope (*r*). Instantaneous bout rate increased linearly with the absolute value of 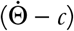; to account for this, two lines were fit to calculate two y-intercepts, one for negative values of 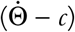 (*b*_down_) and one for positive values (*b*_up_). Baseline bout rate (*β*) was calculated from the mean y-intercept of the two best-fit lines. Lines were then re-fit to the data with y-intercepts fixed at *β* to calculate the slopes of the two best-fit lines (*r*_down_, *r*_up_).

Bout initiation probability was updated over time based on **Θ**(*t*) and 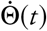, and was also made time-variant, with bout probability dropping to 0 in the first 100 ms (4 time samples) following a bout to match the empirical swim refractory period. After the 100 ms refractory period, swim probability increases to the full bout probability as a function of time elapsed from the last bout (*t* − *t_bout_*):

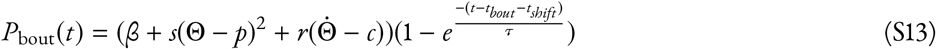

Two parameters determined the shape and rise time of the time-dependency of *P_bout_*: a time shift parameter, *t_shift_* (in samples) and rise-time, *τ* (in samples). For each condition, these parameters were fit to minimize the difference between simulated interbout interval distribution with the empirical distribution.

To test that the modified version of swim simulation described here behaved in a similar manner to our previously published model[37], we compared our full model (described above) using parameters fit from control data with two null models with altered bout initiation and 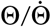 bout correction terms in the model. In the “Bout Timing Null Model,” bouts were initiated randomly in a pitch and angular velocity-independent manner:

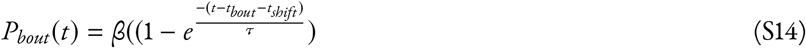

In the “Bout Correction Null Model”, **Θ***_pre_* and 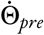 were not correlated with **ΔΘ** or 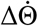, and were instead drawn randomly from a Gaussian distribution with mean and standard deviation matched to empirical **ΔΘ** or 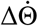 distributions. In the “Complete Null Model” both bout initiation and bout correction terms of the model were randomly drawn. Simulated bouts from the Bout Timing Null Model and Complete Null Model were less well balanced than the Full Model, and performed similarly to our previously published model (6B). To quantify the discriminability between the simulated pitch distributions of our full and null models and empirical pitch distributions from control larvae, we used the area under the receiver operating characteristic (AUROC), where fully discriminable distributions have an AUROC of 1 and identical distributions have an AUROC of 0.5. The comparison of the Full Control model to the empirical control data had an AUROC of 0.48. The Full Control model captured almost the entirety of the variability (86% of empirical standard deviation) seen in the empirical control distribution

To test the effects of vestibulospinal neuron lesions on pitch distributions of simulated larvae, we replaced subsets of parameters in the Full Control model with parameters fit from vestibulospinal lesion empirical data. In the Bout Timing model, *P_bout_*(*t*) was calculated using *β_lesion_, s_lesion_, r_lesion_*, *t_lesionshift_*, and *τ_lesion_*, with all other parameters calculated from control data. In the Bout Correction model, 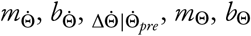, and ΔΘ|Θ_*pre*_ were replaced with parameters fit from lesion data, and all other computations calculated with control parameters. The comparison of the Full Lesion model to the empirical lesion data had an AUROC of 0.44. The Full Lesion Model was more stable than the empirical lesion data, capturing 71% of the postural variability (standard deviation) seen in the empirical vestibulospinal lesioned pitch distributions.

### Statistics

The expected value and variance of data are reported as the mean and the standard deviation or the median and median absolute difference. When data satisfied criteria of normality, parametric statistical tests were used, otherwise we used their nonparametric counterparts. Paired statistics were used for comparisons between siblings. Criteria for significance was set at 0.05 and, when applicable, corrected for multiple comparisons.

### Data sharing

All raw data and code for analysis are available online at http://www.schoppiklab.com/

## Supporting information

Full manuscript with line numbers and bigger margin

## Acknowledgments

Research was supported by the National Institute on Deafness and Communication Disorders or the National Institutes of Health under award numbers DC012775, DC016316, and DC019554. The authors would like to thank Başak Sevinç for assistance with fish maintenance and care, and Katherine Nagel, Niels Ringstad, Mike Long, Dan Sanes, Dora Angelaki along with the members of the Schoppik and Nagel lab for their valuable feedback and discussions.

## Author Contributions

Conceptualization: KRH and DS, Methodology: KRH, KH, MG, and DS, Investigation: KRH, KH, MG, ZD, Visualization: KRH Writing: KRH Editing: DS, Funding Acquisition: KRH and DS, Supervision: DS.

## Author Competing Interests

The authors declare no competing interests.

---

**Figure S1:**
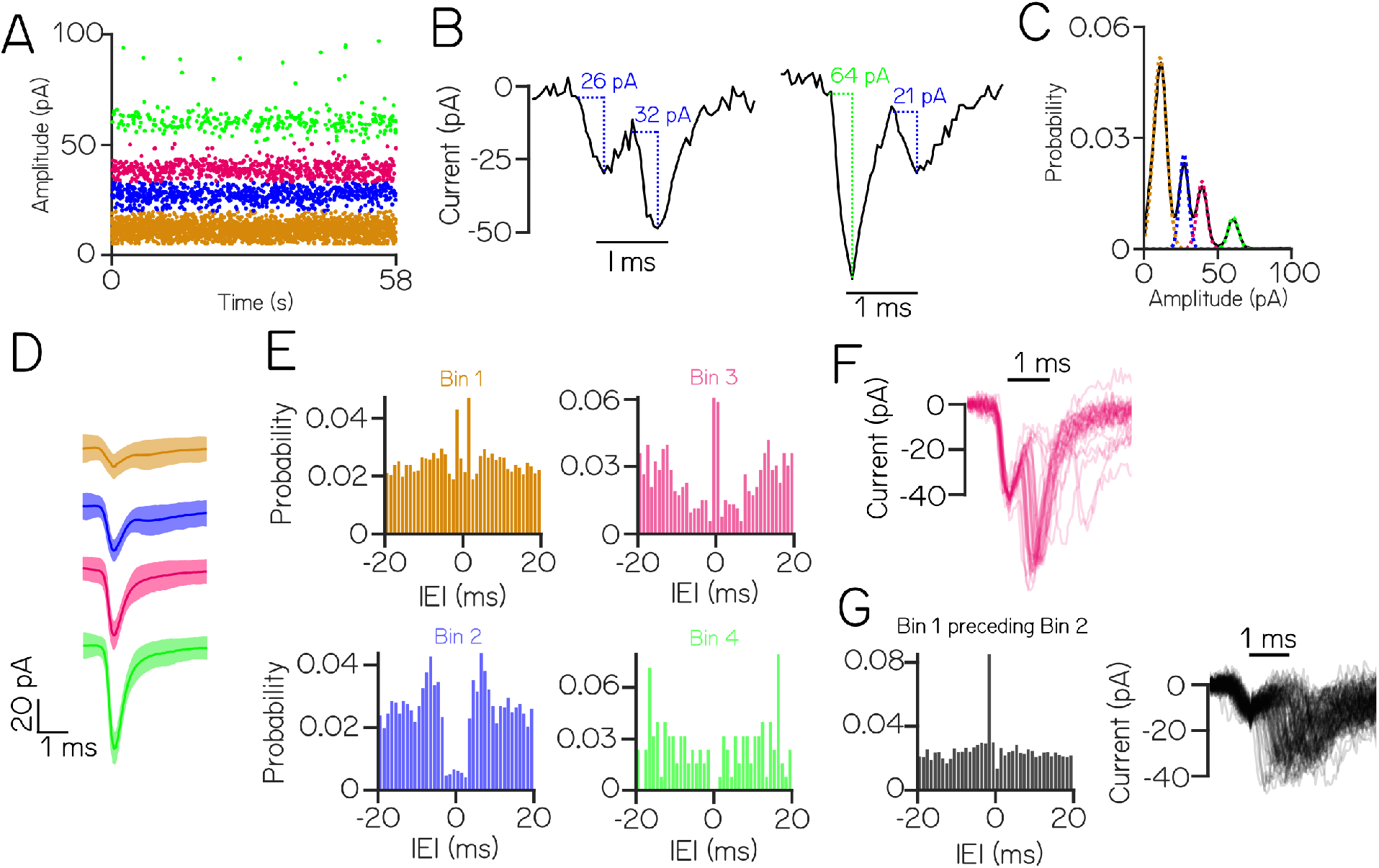
Low-amplitude EPSCs events within the same amplitude bin reflect multiple neuronal inputs. (A) An example cell with four stationary and discrete amplitude bins (color) (B) Example EPSC traces demonstrate events that co-occur within 1 ms. Event pairs are either within-bin (left) or across-bin (right) (C) To estimate an upper limit on the expected refractory period violations due to bin overlap, EPSC amplitude distributions were modeled as a sum of individual Gaussians (colors) (D) Average waveforms from each EPSC bin (± S.D.) (E) Auto-correlograms show structure of inter-event intervals within an EPSC bin; note peaks near zero in bins 1&3, a non-zero valley for bin 2, and a true valley for the high-amplitude bin 4 (F) Waveforms from EPSC pairs within bin 3 with latencies < 2ms (G) Left: cross-correlogram between bin 1 and bin 2, Right: waveforms of EPSC pairs between 1 & 2 with latency < 2ms. The small preceding event and large jitter between peaks are both inconsistent with the expected profile of an electrochemical synapse

**Figure S2:**
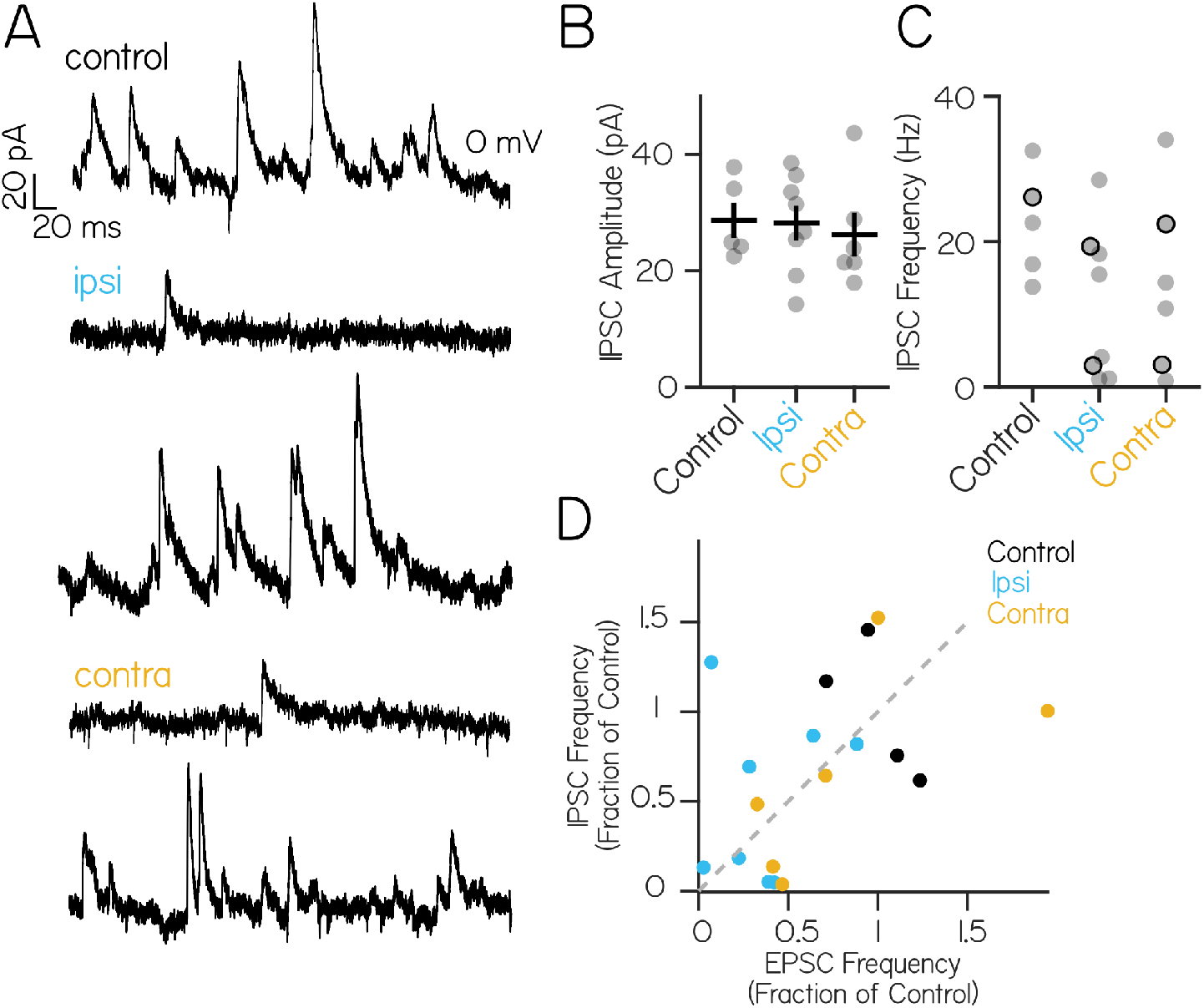
Inhibitory currents onto a single vestibulospinal neuron can sometimes be disrupted by contralateral or ipsilateral utricular lesions; this disruption is not correlated with impairments to excitatory currents. (A) Control trace from a neuron held at 0mV shows inhibitory input at rest (top). After ipsilateral (middle) or contralateral (bottom) otic capsule lesion, some cells experience strong loss of inhibitory currents, while others appear unaffected. (B) IPSC amplitude is unchanged across control and lesion conditions (Kruskal-Wallis H(2)=0.94, p=0.24) (C) The distribution of IPSC frequency after ipsilateral or contralateral lesions is bimodal. Black outlined dots represent cells for example traces in (A). No control neuron (n=5) experienced IPSC frequency less than 10 Hz. After ipsilateral lesions, 4/8 neurons experienced IPSC frequencies less than 10 Hz. After contralateral lesion, 2/6 neurons received IPSCs below 10 Hz (D) Spontaneous EPSC frequency and spontaneous IPSC frequency in recorded neurons from control (black), ipsilateral (blue) and contralateral (yellow) lesion conditions. Each dot is a neuron where both spontaneous excitatory and inhibitory current traces were recorded. For comparison, frequencies were normalized to the mean IPSC or EPSC frequency of control cells. Gray dashes indicate the unity line

**Figure S3:**
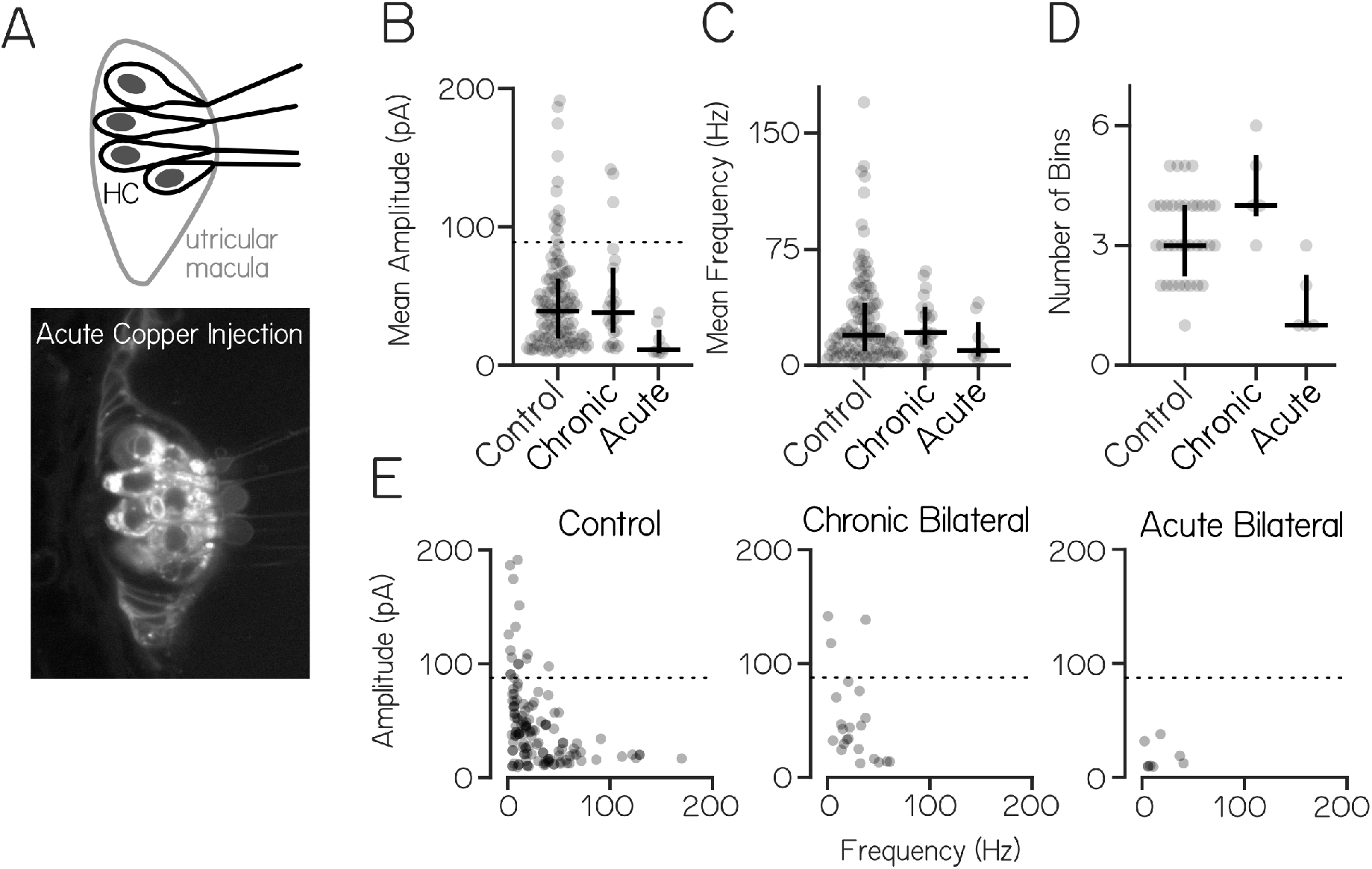
Phasic tuning of action potentials, membrane potential, and synaptic events during linear translation. (A) Phase of peak firing response to lateral and fore-aft stimuli. The majority of lateral-responsive cells respond to contralateral translation (6/10), and the majority of fore-aft responsive cells responded to rostral translation (7/10) (B) Representative low-pass filtered voltage traces during lateral translation from control (black), chronic (purple), and acute lesion (orange) shows rhythmic oscillation (solid) and sinusoidal fits (dashed) of membrane potential which are attenuated after lesion (C) During lateral translation, slow frequency voltage modulation is high in cells from control fish (3.4 mV, ± 0.5 SEM lateral, n=17 cells). Amplitude modulation is reduced after chronic (0.2 mV ± 0.2 SEM lateral, n=4 cells; p=0.02) but not acute utricle lesions (1. mV ± 0.7 SEM lateral, n=4 cells; p=0.16) (D) Slow frequency voltage modulation amplitude is positively correlated with the modulation in the mean EPSC event rate in both lateral (Pearson’s r=0.59 (R^2^=0.36) and fore-aft (Pearson’s r=0.67 (R^2^=0.45) translation (E-F) IPSCs and EPSCs from two cells show out-of-phase (Cell 1) or similar (Cell 2) responses to lateral translation. Top: acceleration with timing of peak response peak overlaid (blue/pink dashed line). Middle: spike rasters bottom: PSTH (F) Inhibitory and excitatory inputs from another example cell are cotuned to lateral acceleration, with peak IPSC and EPSC rates occurring in-phase with ipsilateral acceleration G) Phase comparison of EPSC and IPSC peak rate of two example cells.

**Figure S4:**
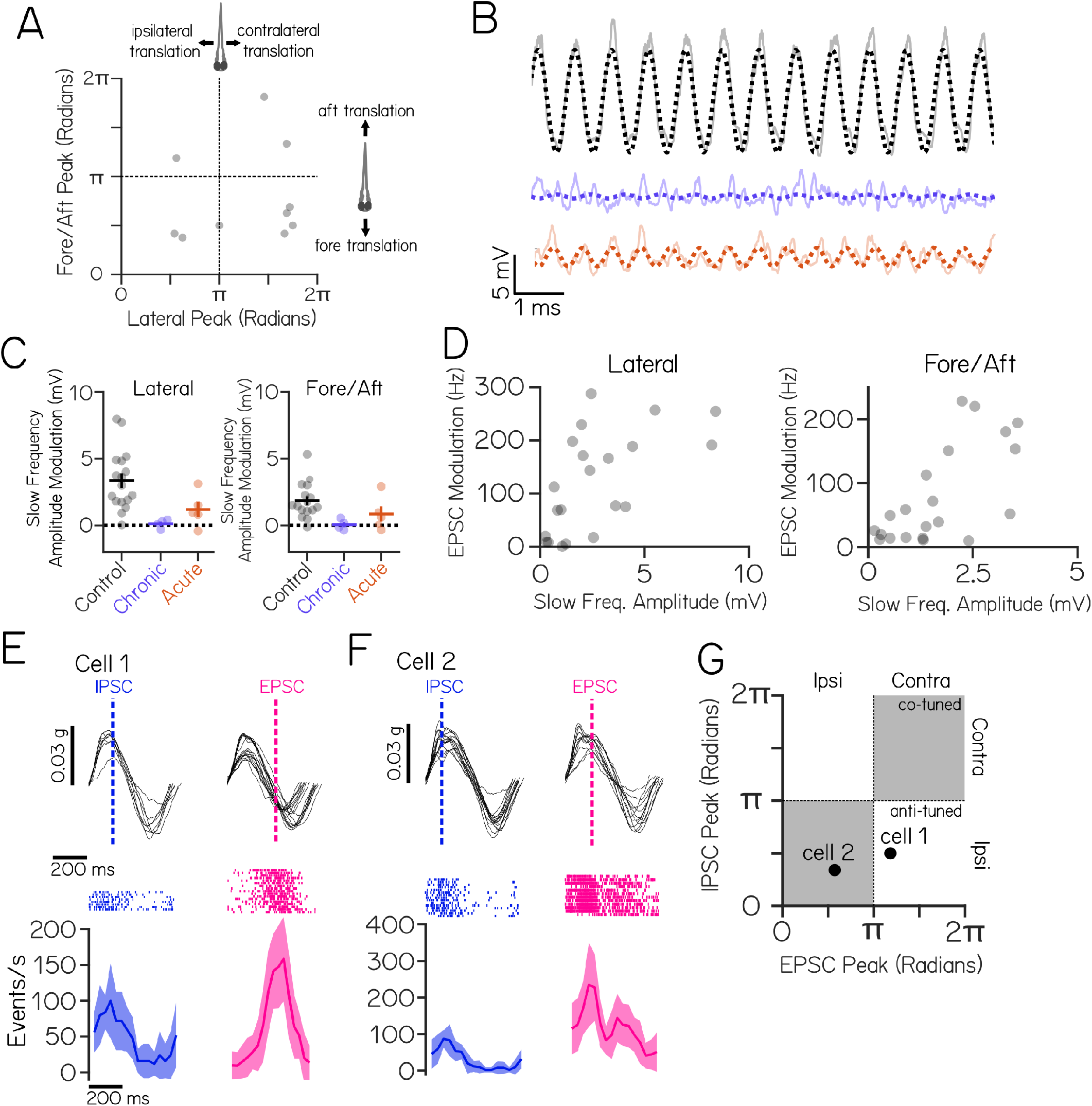
Spontaneous excitatory synaptic input persists after chronic bilateral utricle loss (*otogelin* mutant) but is nearly abolished after acute bilateral lesions (chemical lesion of hair cells). (A) Top: schematic of hair cells within the utricular macula of the inner ear. Bottom: disrupted hair cell membranes after copper injection (B) Mean amplitude of EPSC bins (gray circles) from vestibulospinal neurons in control, chronic and acute conditions. Median and inter-quartile range of data. “High amplitude” bins (top 5% of control) above black dashed line (C) Mean event frequency of EPSC bins. Median and inter-quartile range of data (D) Number of EPSC bins in a cell. Each dot is one vestibulospinal cell. Median and inter-quartile range of data (E) Mean amplitude of an EPSC bin (gray circle) as a function of frequency in control, chronic, and acute conditions. Dashed line delineates “high amplitude” bins

**Figure S5:**
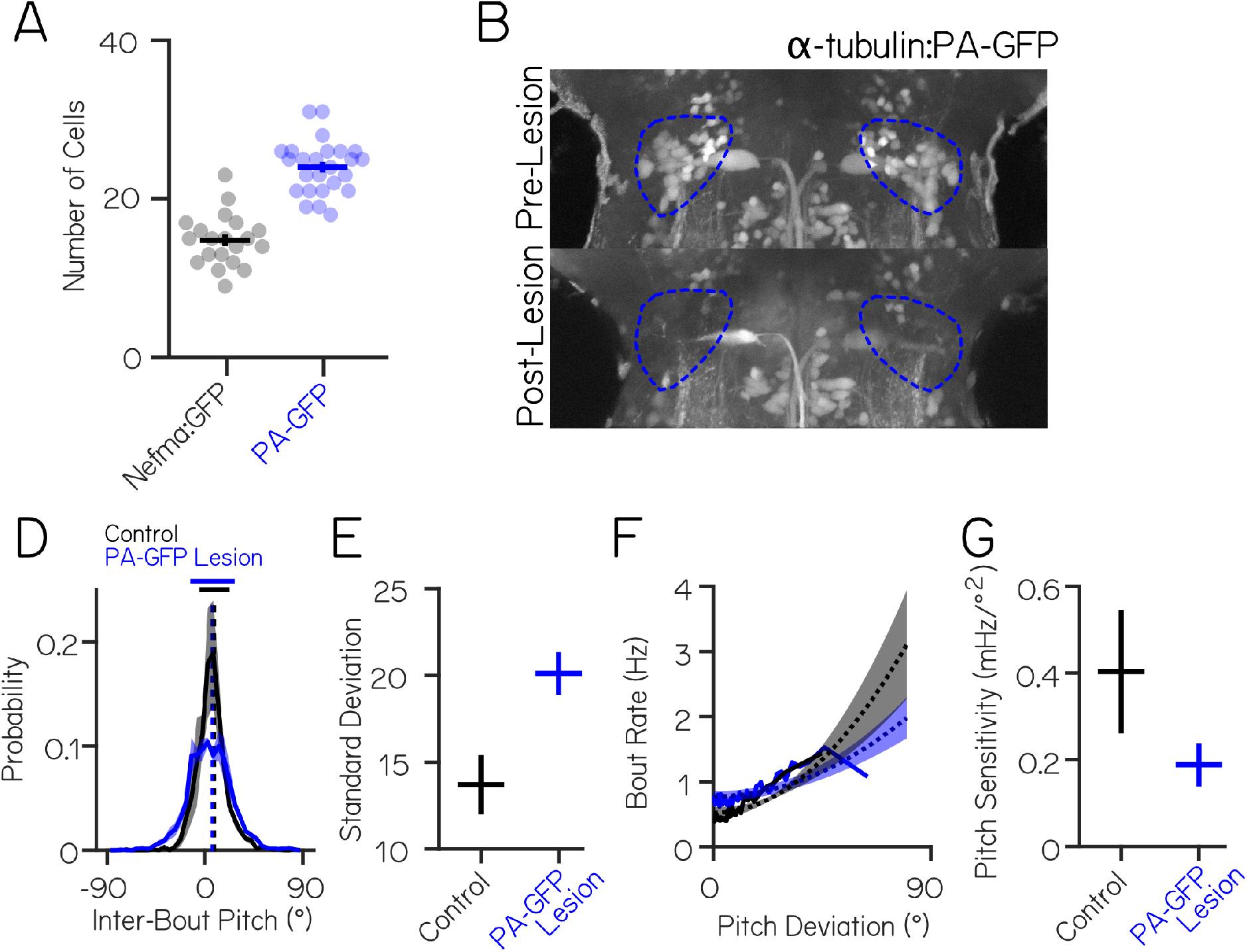
Lesions of a larger pool of optically identified vestibulospinal neurons replicates postural and coordination disruption observed in *Tg(nefma:GAL4);Tg(UAS:GFP)* lesions. (A) *Tg(nefma:GAL4);Tg(UAS:GFP)* labels fewer (14.8 ± 0.7 SEM, n=20 hemispheres, 10 fish) neurons than optically-backfilled *Tg(α-tubulin:C3PA-GFP)* fish (24 (± 0.6 SEM, n=25 hemispheres, 13 fish, unpaired t-test p=8.3*10^−12^) (B) Dorsoventral distribution of vestibulospinal cells in both lines. *Tg(nefma:GAL4);Tg(UAS:GFP)* labels fewer cells in the dorsal vestibulospinal nucleus compared to photofills in *Tg(α-tubulin:C3PA-GFP)* (C) Representative maximum intensity projection of spinal projecting neurons in the hindbrain of a *Tg(α-tubulin:C3PA-GFP)* fish following optical backfill before (top) and after (bottom) two-photon mediated photoablation (D) Average probability distributions (± S.D) of inter-bout pitch angle for sibling controls (black, n=17 fish, 1828 bouts control) and vestibulospinal lesioned fish (blue, n=17 fish, 2125 bouts lesion) show no change in average posture (dashed vertical lines) but greater variability (solid horizontal lines ± 1 S.D.) (E) Standard deviation of pitch is higher in vestibulospinal lesioned fish (20.1±1.2) compared to sibling controls (13.7 ±1.7, paired t-test p=1.0*10^−5^) (F) Bout rate as a function of deviation from preferred posture for lesions (blue) and control siblings (black). Solid lines represent raw data, dashed lines represent parabolic fits to raw data ± S.D. of the fit estimates. (G) Pitch sensitivity (parabolic fit) is decreased in vestibulospinal lesioned fish (0.2 ± 0.05 S.D., paired t-test p=0.002) compared to sibling controls (0.4 ± 0.14 S.D.)

## Notes

### Competing Interest Statement

The authors have declared no competing interest.

