## Supplementary material for "Synaptic Encoding of Vestibular Sensation Regulates Movement Timing and Coordination": Full manuscript with line numbers and bigger margin

$$\phi_{\mathrm{A}} = \beta_{\mathrm{A}}(29 \cdot H_{\mathrm{A}} \cdot F_{\mathrm{A}}) \tag{S2}$$

where $\beta_{\mathrm{A}}$ is the number of observed EPSC events falling with amplitude bin, $A$ (defined by EPSC amplitude thresholds from $a$ to $b$ pA). The hit event rate (events assigned to bin $A$ that truly derived from bin $A$) was calculated by $H_{\mathrm{A}}$:

$$H_{\mathrm{A}} = \frac{\beta \cdot \int_a^b p(I_{\mathrm{A}})dI}{s} \tag{S3}$$

where $\beta$ is the total number of observed EPSCs events across all amplitude bins, and $s$ is the trial length in samples. The false-positive event rate for bin $A$ (events falsely assigned to bin $A$ when they derived from the overlapping tails of Gaussian distributions from other bins) was calculated by $F_{\mathrm{A}}$:

$$F_{\mathrm{A}} = \frac{\beta \cdot (\int_a^b p(I_{\mathrm{B}})dI + \int_a^b p(I_{\mathrm{C}})dI + ... + \int_a^b p(I_{\mathrm{N}})dI}{s} \tag{S4}$$

where $p(I_{\mathrm{B}})$ is the estimated probability distribution of the second EPSC amplitude bin, and $p(I_{\mathrm{N}})$ is the $\mathrm{n}^{\mathrm{th}}$ EPSC amplitude bin.

As described previously [39], a logistic function was used to fit the relationship between attack angle ($\gamma$) and posture change($r$), based on the formula

$$\gamma(r) = \gamma_0 + \frac{\gamma_{max}}{1 + e^{-k(r-r_0)}} \tag{S7}$$

where $\gamma_0$ gives the most negative attack angle (in deg), ($\gamma_{max} + \gamma_0$) gives that largest positive attack angle (in deg), and $k$ is the steepness parameter (in deg$^{-1}$). From the derivative of S7, sigmoid maximal slope (found at $r$=$r_0$) is given by $k\gamma_{max}/4$. Sigmoid center position ($r_0$) was itself defined from a parameter for rise position ($r_{rise}$, posture change at which the sigmoid rises to 1/8 of its upper asymptote:

$$r_0 = \frac{kr_{rise} + log(\frac{-\gamma_0 - 7\gamma_{max}}{\gamma_0 + \gamma_{max}})}{k} \tag{S8}$$

Parameter fits were estimated in Matlab using a nonlinear regression based solver (trust-region-reflective). Initial parameter values were $k$=1 deg$^{-1}$, $\gamma_0$=-0.2°, $\gamma_{max}$=10°, and $r_{rise}$=-1°. As previously [39], only data from swims that occurred during circadian day (as determined by incubator light cycle) were included. To improve fit, we excluded attack angles > | ± 50°| (1.3% and 1.6% of observations in lesions and controls respectively). To compare sigmoid slope between conditions, $\gamma_0$, $\gamma_{\text{max}}$, and $r_{\text{rise}}$ were fixed at means between lesion and control conditions to fit a one-parameter sigmoid.

Due to low bout numbers per experiment (443 ± 271 bouts per clutch for *Tg(nefma:GAL4);Tg(UAS:GFP)* experiments), a single experiment often did not contain enough data to sufficiently sample the entire distribution of postures. Therefore data for all calculations were pooled across all bouts in a given group (lesion or control). Estimates of variation across clutches was performed using jackknifed resampling, where each sample left out two experimental clutch replicates. Paired statistics for behavioral measurements used paired jackknifed sub-samples.

### Behavioral Modeling

We generated condition-specific swimming simulations using a generative swim model described previously [37], with updates to the model to improve the fit to the empirical control dataset collected here. In each condition, 20 simulated fish swam for 3000 seconds with discrete time steps equivalent to those in the captured data ($\Delta t = 25$ ms). At each time step ($t$), the pitch ($\Theta(t)$) is updated due to passive posture destabilization from angular acceleration, with $\Theta(t)$ initialized at a randomly drawn integer from a uniform distribution between ±90°. Angular velocity ($\dot{\Theta}$) was initialized at 0, and was calculated as the sum of $\dot{\Theta}(t-1)$ and the integral of angular acceleration, $\ddot{\Theta}(t)$:

$$\Theta(t) = \Theta(t-1) + \dot{\Theta}(t)\Delta t \tag{S9}$$

Between swim bouts, simulated larvae were destabilized according to angular acceleration. Angular acceleration varied as a function of pitch in the preceding time step ($\Theta(t-1)$:

$$\ddot{\Theta}(t) = \ddot{\Theta}_c cos(\Theta(t-1)) \tag{S10}$$

where $\ddot{\Theta}_c$ is the maximal empirical angular acceleration observed between bouts for fish each condition, calculated by taking the median of the second differential of smoothed pitch angles between bouts using a moving average filter with a span of 75 ms.

Simulated larval pitch was updated across time according to passive destabilization until a swim bout was initiated. When a bout was initiated, pitch and angular velocity were updated according to condition-specific correction parameters based on empirical swim bout kinematics. Angular velocity correction from swim bouts was corrected by making net angular acceleration across swim bouts ($\Delta\dot{\Theta}$) correlated with pre-bout angular velocity ($\dot{\Theta}_{\text{pre}}$). Condition-specific correlations were determined by best-fit lines to empirical data, defined by a slope ($m_{\dot{\Theta}}$)and intercept ($b_{\dot{\Theta}}$) for both positive (nose-up) and negative (nose-down) angular velocities.

To reproduce the empirical variability of bout kinematics, net angular acceleration incorporated a noise term ($\varepsilon_{\Delta\dot{\Theta}|\ddot{\Theta}}$) drawn randomly from a Gaussian distribution with mean of 0, and standard deviation ($\sigma_{\Delta\dot{\Theta}|\ddot{\Theta}}$) calculated from the empirical standard deviation of $\Delta\dot{\vartheta}$ and reduced proportionally by the unexplained variability from the correlation between $\Delta\dot{\Theta}$ and $\dot{\Theta}_{\text{pre}}$.

A bout initiated at time $t$ corrected $\dot{\Theta}$ after completion of the bout, 100 ms later (4 time samples, matched to empirical bout duration):

$$\dot{\Theta}(t+4) = \dot{\Theta}(t) + m_{\dot{\Theta}}\dot{\Theta}(t) + b_{\dot{\Theta}} + \varepsilon_{\Delta\dot{\Theta}|\dot{\Theta}_{\text{pre}}} \tag{S11}$$

The same approach was used to condition net bout rotation ($\Delta\Theta$) on pre-bout pitch ($\Theta_{\text{pre}}$) based on a single best-fit line, with a single slope ($m_{\Theta}$ and intercept ($b_{\Theta}$). The corresponding noise term ($\varepsilon_{\Theta}$) was not reduced by the unexplained variability to better match empirical pitch distributions. For all conditions, the fit line was constrained to the linear portion of the relationship (-30 to +40° pre-bout pitch). If the simulated fish's posture prior to a bout was outside of this window (only 1% of simulated inter-bouts), net bout rotation was calculated using $m_{-30}$, $b_{-30}$ if $\Theta < -30$, or $m_{40}$, $b_{40}$ if $\Theta > 40$. To further impose a ceiling on the counter-rotation values attainable by our simulated larvae, a maximum net bout rotation was imposed ($\Delta\Theta_{\max}$=±26.3°) based on empirical values; if $\Delta\Theta > \Delta\Theta_{\max}$, then $\Delta\Theta$ was set to $\Delta\Theta_{\max}$.

Bouts occurred in the model based an internal state variable representing the probability of bout initiation ($P_{\text{bout}}$). Bout initiation was calculated as the sum of a non-posture dependent baseline bout rate variable ($\beta$), a $\Theta$-dependent bout rate, and $\dot{\Theta}$-dependent bout rate:

$$P_{\text{bout}} = \beta + s(\Theta - p)^2 + r(\dot{\Theta} - c) \tag{S12}$$

Bout rate as a function of pitch was calculated by fitting a parabola to the relationship between instantaneous bout rate and deviation from mean posture ($\Theta - p$) to get a pitch sensitivity parameter ($s$) for each condition. Bout rate as a function of angular velocity was calculated by fitting a line to the correlation between instantaneous bout rate and mean-subtracted angular velocity ($\dot{\Theta} - c$) to get a slope ($r$). Instantaneous bout rate increased linearly with the absolute value of ($\dot{\Theta} - c$); to account for this, two lines were fit to calculate two y-intercepts, one for negative values of ($\dot{\Theta} - c$) ($b_{\text{down}}$) and one for positive values ($b_{\text{up}}$). Baseline bout rate ($\beta$) was calculated from the mean y-intercept of the two best-fit lines. Lines were then re-fit to the data with y-intercepts fixed at $\beta$ to calculate the slopes of the two best-fit lines ($r_{\text{down}}$, $r_{\text{up}}$).

Bout initiation probability was updated over time based on $\Theta(t)$ and $\dot{\Theta}(t)$, and was also made time-variant, with bout probability dropping to 0 in the first 100 ms (4 time samples) following a bout to match the empirical swim refractory period. After the 100 ms refractory period, swim probability increases to the full bout probability as a function of time elapsed from the last bout ($t - t_{bout}$):

$$P_{\text{bout}}(t) = (\beta + s(\Theta - p)^2 + r(\dot{\Theta} - c))(1 - e^{\frac{-(t - t_{bout} - t_{shift})}{\tau}}) \tag{S13}$$

Two parameters determined the shape and rise time of the time-dependency of $P_{bout}$: a time shift parameter, $t_{shift}$ (in samples) and rise-time, $\tau$ (in samples). For each condition, these parameters were fit to minimize the difference between simulated inter-bout interval distribution with the empirical distribution.

To test that the modified version of swim simulation described here behaved in a similar manner to our previously published model[37], we compared our full model (described above) using parameters fit from control data with two null models with altered bout initiation and $\Theta/\dot{\Theta}$ bout correction terms in the model. In the "Bout Timing Null Model," bouts were initiated randomly in a pitch and angular velocity-independent manner:

$$P_{bout}(t) = \beta((1 - e^{\frac{-(t - t_{bout} - t_{shift})}{\tau}}) \tag{S14}$$

In the "Bout Correction Null Model", $\Theta_{pre}$ and $\dot{\Theta}_{pre}$ were not correlated with $\Delta\Theta$ or $\Delta\dot{\Theta}$, and were instead drawn randomly from a Gaussian distribution with mean and standard deviation matched to empirical $\Delta\Theta$ or $\Delta\dot{\Theta}$ distributions. In the "Complete Null Model" both bout initiation and bout correction terms of the model were randomly drawn. Simulated bouts from the Bout Timing Null Model and Complete Null Model were less well balanced than the Full Model, and performed similarly to our previously published model (6B).

To quantify the discriminability between the simulated pitch distributions of our full and null models and empirical pitch distributions from control larvae, we used the area under the receiver operating characteristic (AUROC), where fully discriminable distributions have an AUROC of 1 and identical distributions have an AUROC of 0.5. The comparison of the Full Control model to the empirical control data had an AUROC of 0.48. The Full Control model captured almost the entirety of the variability (86% of empirical standard deviation) seen in the empirical control distribution

To test the effects of vestibulospinal neuron lesions on pitch distributions of simulated larvae, we replaced subsets of parameters in the Full Control model with parameters fit from vestibulospinal lesion empirical data. In the Bout Timing model, $P_{bout}(t)$ was calculated using $\beta_{lesion}$, $s_{lesion}$, $r_{lesion}$, $t_{lesionshift}$, and $\tau_{lesion}$, with all other parameters calculated from control data. In the Bout Correction model, $m_{\dot{\Theta}}$, $b_{\dot{\Theta}}$, $_{\Delta\dot{\Theta}|\dot{\Theta}_{pre}}$, $m_{\Theta}$, $b_{\Theta}$, and $_{\Delta\Theta|\Theta_{pre}}$

were replaced with parameters fit from lesion data, and all other computations calculated with control parameters. The comparison of the Full Lesion model to the empirical lesion data had an AUROC of 0.44. The Full Lesion Model was more stable than the empirical lesion data, capturing 71% of the postural variability (standard deviation) seen in the empirical vestibulospinal lesioned pitch distributions.

#### Statistics

The expected value and variance of data are reported as the mean and the standard deviation or the median and median absolute difference. When data satisfied criteria of normality, parametric statistical tests were used, otherwise we used their non-parametric counterparts. Paired statistics were used for comparisons between siblings. Criteria for significance was set at 0.05 and, when applicable, corrected for multiple comparisons.

#### Data sharing

All raw data and code for analysis are available online at http://www.schoppiklab.com/

### Acknowledgments

Research was supported by the National Institute on Deafness and Communication Disorders or the National Institutes of Health under award numbers DC012775, DC016316, and DC019554. The authors would like to thank Başak Sevinç for assistance with fish maintenance and care, and Katherine Nagel, Niels Ringstad, Mike Long, Dan Sanes, Dora Angelaki along with the members of the Schoppik and Nagel lab for their valuable feedback and discussions.

### Author Contributions

Conceptualization: KRH and DS, Methodology: KRH, KH, MG, and DS, Investigation: KRH, KH, MG, ZD, Visualization: KRH Writing: KRH Editing: DS, Funding Acquisition: KRH and DS, Supervision: DS.

### Author Competing Interests

The authors declare no competing interests.

Table 1: Spontaneous properties across conditions

| | Unit | Control | Cs Control | Ipsi Lesion | Contra Lesion | Bilateral Lesion | Rock Solo |
|---|---|---|---|---|---|---|---|
| Resting Membrane Potential | mV | -67.3 | -60.8 | -63.7 | -63.8 | -68.9 | -66 |
| Series Resistance | MΩ | 34.7 | 25.8 | 43.6 | 45.7 | 38.4 | 31.2 |
| Input Resistance | MΩ | 236 | 202 | 264 | 178 | 310 | 228 |
| Resting Firing Frequency | Hz | 1.4 | - | - | - | 0 | 0 |
| Rheobase | pA | 111.8 | - | - | - | 38.9 | 58.8 |
| Resting EPSC Frequency | Hz | 101.3 | 65.4 | 30.2 | 59.2 | 46.4 | 119.2 |

Table 2: Behavioral properties

| Variable | Unit | Nefma::GFP | Nefma::GFP lesioned | PA-GFP | PA-GFP lesioned |
|---|---|---|---|---|---|
| Preferred posture (±S.D) | deg | 9.8 (±15.6) | 10.4 (±23.3) | 8.6 (±13.7) | 6.8(± 20.1) |
| Inter-bout interval (± M.A.D) | s | 1.7 (±2.6) | 1.6 (±2.5) | 2.3 (±1.5) | 1.3 (±1.2) |
| Bout duration (± M.A.D) | s | 0.10 (±0.06) | 0.13 (±0.06) | 0.13 (±0.04) | 0.13 (±0.04) |
| Bout displacement (± M.A.D) | mm | 1.1 (±0.7) | 1.2 (±0.7) | 1.3 (±0.6) | 1.4 (±0.6) |
| Maximum linear speed (± M.A.D) | mm/s | 9.6 (±3.8) | 10.0 (±3.6) | 11.2 (±4.3) | 12.9 (±4.4) |

Table 3: Modeling parameters

| Parameter | Symbol | Unit | Nefma::GFP | Nefma::GFP lesioned |
|---|---|---|---|---|
| Pitch sensitivity | $s$ | mHz/deg$^2$ | 0.30 | 0.12 |
| Angular velocity sensitivity (nose-down) | $r_{down}$ | deg$^{-1}$ | -0.12 | -0.09 |
| Angular velocity sensitivity (nose-up) | $r_{up}$ | deg$^{-1}$ | 0.13 | 0.09 |
| Mean posture | $p$ | deg | 9.8 | 10.4 |
| Median angular velocity | $c$ | deg/s | -0.15 | -0.66 |
| Baseline bout rate | $\beta$ | Hz | 0.64 | 0.67 |
| Angular Velocity correction gain (nose-down) | $m_{down\dot{\Theta}}$ | - | -0.94 | -0.83 |
| Angular Velocity correction gain (nose-up) | $m_{up\dot{\Theta}}$ | - | -0.99 | -0.98 |
| Pitch correction gain | $m_{\Theta}$ | - | -0.31 | -0.28 |
| Maximal angular acceleration | $\ddot{\Theta}_c$ | deg/s$^2$ | -1.04 | -0.98 |
| Time-dependence shift | $t_{shift}$ | samples | -1.4E5 | -4.9E3 |
| Rise-time | $\tau$ | samples | 1.9E5 | -5.9E3 |

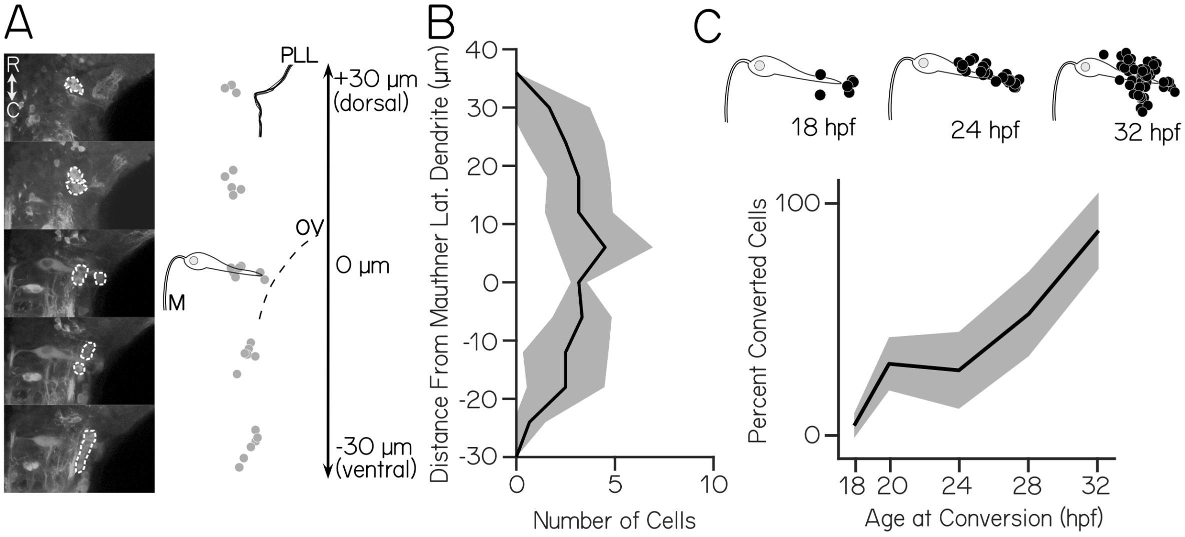


Figure 1: **Vestibulospinal neurons are born well before larvae swim or stabilize posture.** (A) Left: Images of optically backfilled spinal projecting neurons (dashed white) in five dorsoventral planes covering ± 30 um around the Mauthner (M) lateral dendrite, somata. Right: representative neuron location (gray circles) in one hemisphere and the Mauthner lateral dendrite (B) Mean dorsoventral distribution of neurons relative to the Mauthner lateral dendrite per hemisphere (± 1 S.D.) (n=6 fish) (C) Top: neurons (black circles) born at 18, 24, and 32 hpf relative to the Mauthner cell. Bottom: Percent of vestibulospinal neurons born increases steadily to near 100% between 18 and 32 hpf (mean ± 1 S.D.)

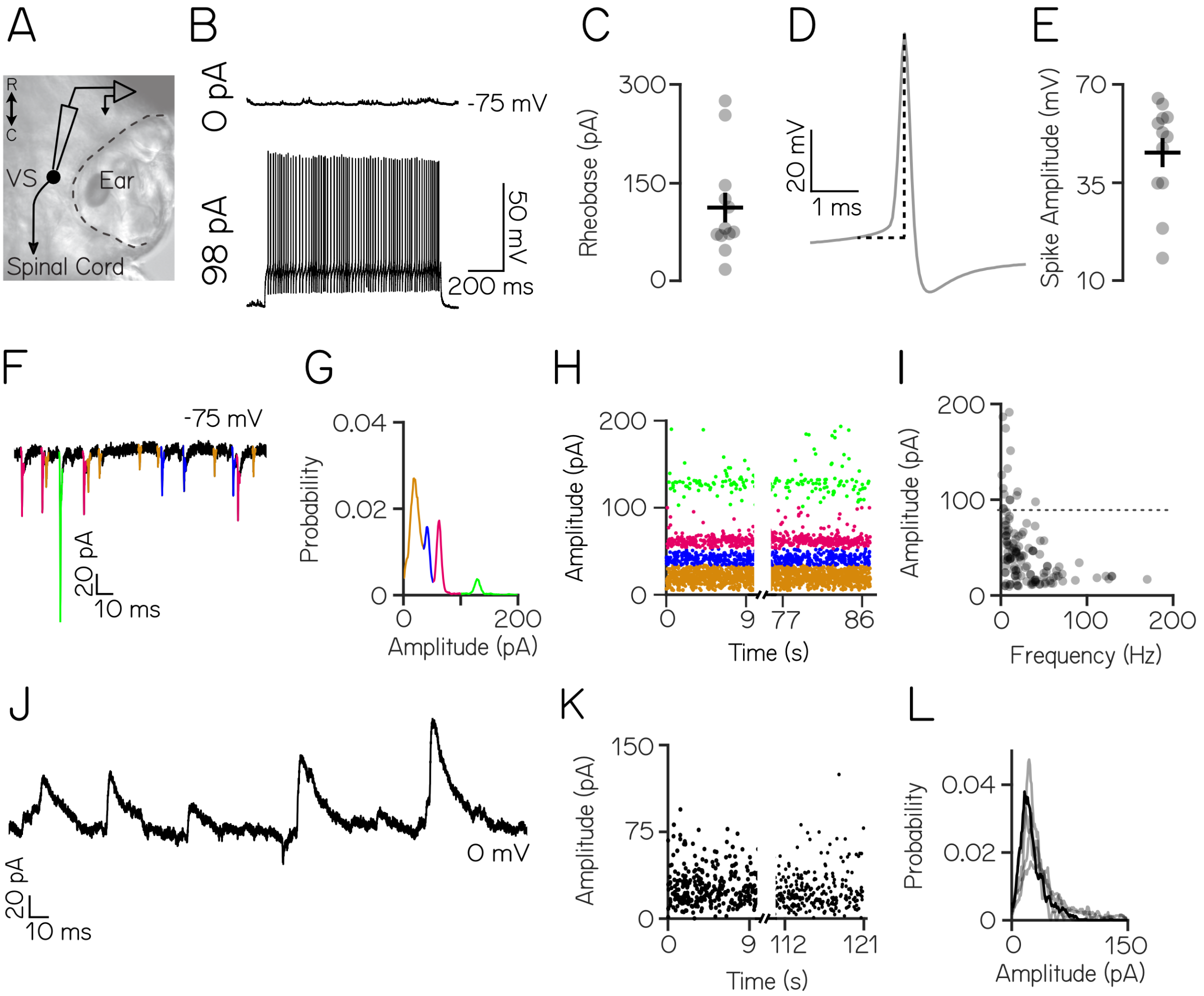


Figure 2: **Larval zebrafish vestibulospinal neurons display mature intrinsic and synaptic properties.** (A) Schematic of spinal-projecting vestibulospinal (VS) neuron targeted for electrophysiology (B) Vestibulospinal response to current injection (98 pA) (C) Rheobase (±SEM) across 21 cells (D) Example action potential waveform with amplitude (dotted line) (E) Action potential average amplitude (±SEM) (F) Distinct amplitudes (color) in excitatory post-synaptic spontaneous currents (EPSCs) from a neuron held at -75mV (G) EPSC amplitudes from a single vestibulospinal cell show 4 distinct probability peaks, or "bins" (color) (H) EPSC bins are stationary in time (I) EPSC amplitude as a function of frequency for all bins in all vestibulospinal cells (116 bins, from 36 cells), dashed line differentiates the top 5% ("high amplitude") of bins (J) Representative current trace from a vestibulospinal neuron held at 0 mV (K) Inhibitory post-synaptic current (IPSC) amplitudes over time (L) IPSC amplitudes for the example neuron in J/K (black line) and other neurons (gray lines) do not show multiple peaks (n=5)

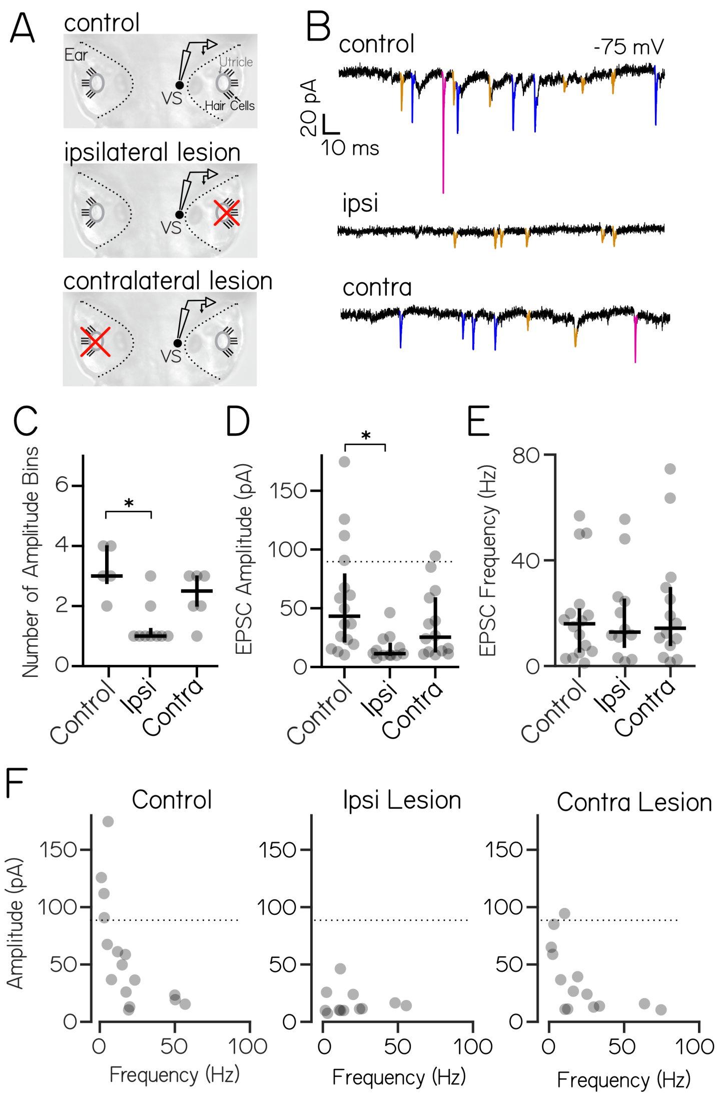


Figure 3: **High amplitude spontaneous excitatory inputs originate in the ipsilateral ear.** (A) Lesion schematic: The utricle (gray circle) was physically removed (red "x") either ipsilateral or contralateral to the recorded vestibulospinal neuron (black circle, "VS") (B) Example current traces from neurons held at -75 mV from control (top), ipsilateral (middle), and contralateral (bottom) experiments; EPSCs in color (C) Number of EPSC amplitude bins per cell (median ± IQR in black) is decreased after ipsilateral, but not contralateral lesion (D) EPSC bin amplitudes (gray circles, median ± IQR in black) are decreased after ipsilateral, but not contralateral, lesion. Dashed line at the top 5% ("high amplitude") of control EPSCs (E) Frequency of events in EPSC bins (gray circles, median ± IQR in black) is unchanged after ipsilateral or contralateral lesion compared to control cells (F) EPSC amplitude vs. frequency for each bin (gray circles) in control and after ipsilateral/contralateral lesions. High amplitude (top 5% of control) bins are lost after ipsilateral lesion

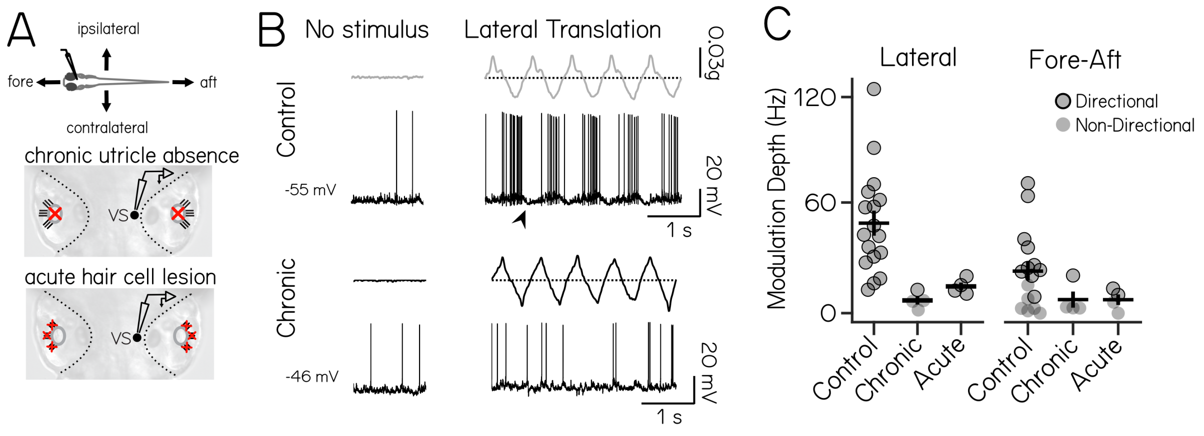


Figure 4: **Vestibulospinal neurons encode utricle-derived body translation.** (A) Immobilized fish were manually translated in the fore-aft or lateral axes (top). Vestibulospinal neurons were recorded in control and after two manipulations: first, in *otogelin* mutants (middle) that do not develop utricles (red "x") and second, after chemically-induced hair cell (red "x") death (bottom) (B) Accelerometer (gray) and voltage trace (black) from a neuron in a control fish (top) showing action potentials and low-amplitude membrane potential fluctuations (arrowhead) in phase with translation. In contrast, activity from a *otogelin* mutant is unaligned with translation (C) Modulation depth of spiking response is disrupted in both the lateral (left) and fore-aft (right) direction after both chronic and acute disruption of the utricle. Gray circles are neurons, black outline denotes statistically significant directional responses

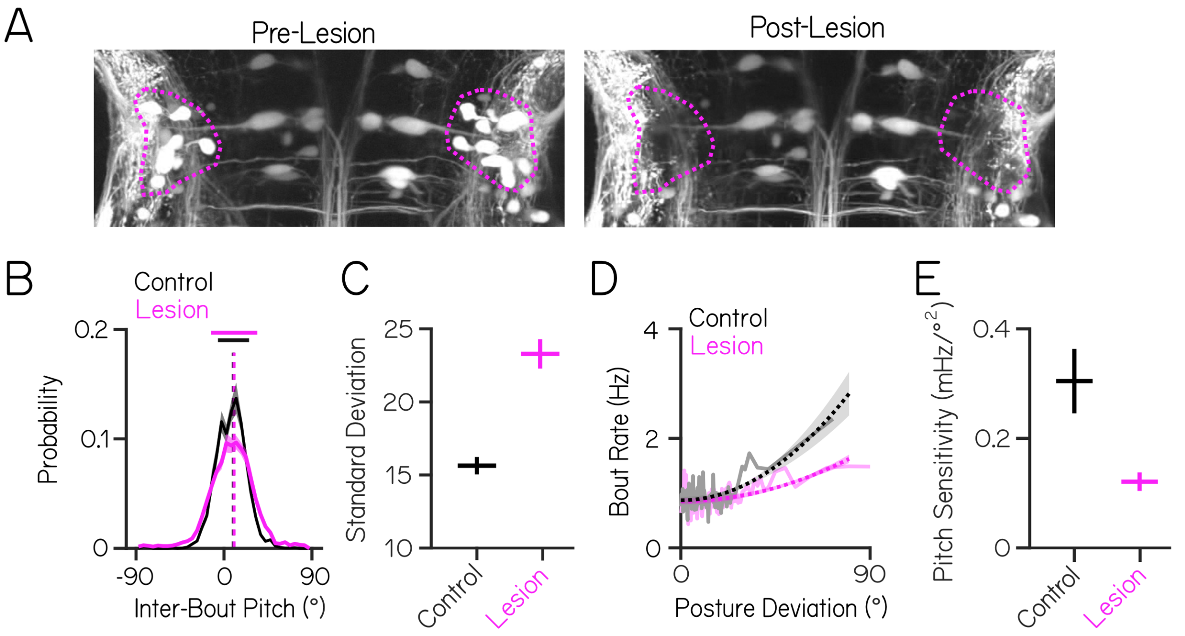


Figure 5: **Vestibulospinal neurons contribute to postural stability and movement timing, but not postural set point.** (A) Images before and after targeted photoablations of genetically-labeled vestibulospinal neurons show selective loss of fluorescent somata in outlined area (magenta) (B) Distributions of observed pitch show no change to average posture (dashed vertical lines) but greater variability (solid horizontal lines ± 1 SD) between control fish (black) and lesioned siblings (magenta) (C) Postural variability (standard deviation of pitch) is greater in lesioned fish than sibling controls; lines are mean ± jacknifed standard deviation (D) Movement timing (bout rate as a function of deviation from preferred posture) in lesioned fish (magenta) and sibling controls (black). Solid lines are binned raw data, dashed lines are parabolic fits to raw data ± 1 SD (E) Pitch sensitivity (parabolic steepness between posture deviation and bout rate) is lower in lesioned fish than sibling controls)

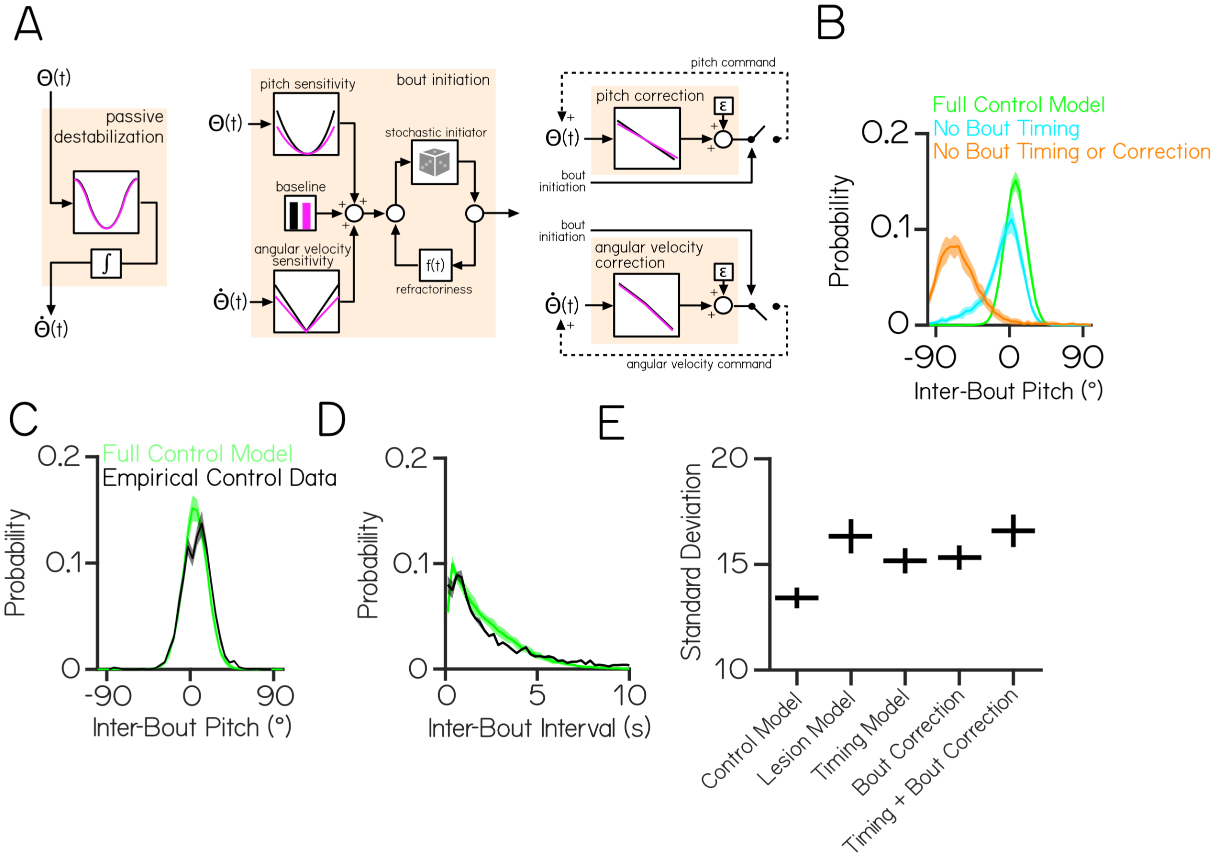


Figure 6: **Behavioral modeling shows that increased postural variability following lesions emerges from combined impairments to swim timing and corrective capacity.** (A) A generative model of swimming adapted from previously published work [37], consists of four computations (tan boxes) determining swim timing, pitch angle, and angular velocity. Each computation contains one or more condition-specific parameters, shown by plots with a black (control) or magenta (vestibulospinal lesion) line (B) Probability distributions of simulated pitch angles in full control model (green), bout timing null model (cyan), and full null model (orange) (C) Probability distributions of observed pitch angles between swim bouts and (D) inter-bout intervals for empirical control data (black) and simulated control fish (green) (E) Standard deviation of simulated pitch probability distributions ±S.D. using different combinations of condition-specific parameters in the computations shown in (A)

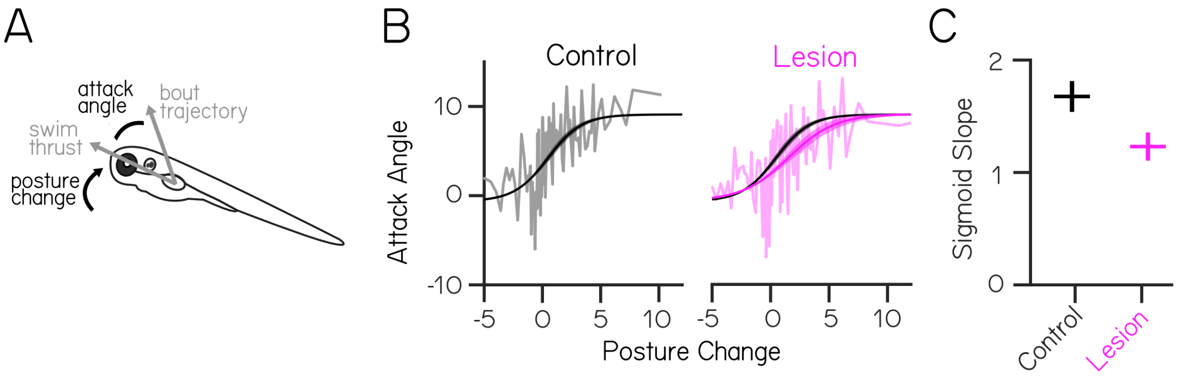


Figure 7: **Vestibulospinal neurons are necessary for coordination during vertical climbs.** (A) Schematic illustrating fin/body coordination during climbing. Larvae climb by coordinating axial rotation ("posture change") and forward thrust with fin movements. The fin contribution can be derived from the attack angle, or the difference between the thrust vector and observed bout trajectory (B) Attack angle as a function of posture change in lesioned fish (magenta) and sibling controls (black). Solid lines are binned raw data, dashed lines represent sigmoidal fits ± 1 SD (C) Sigmoid slope decreases in lesioned fish

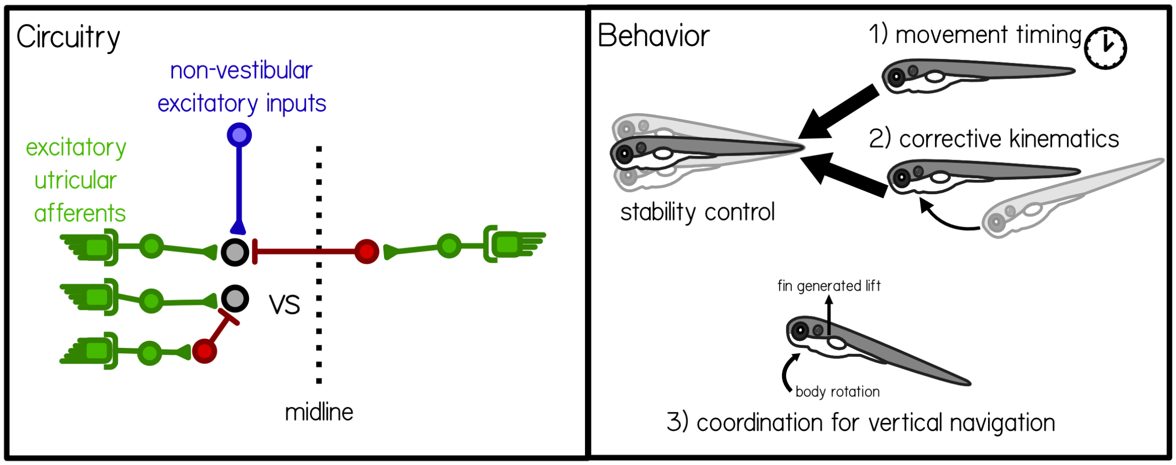


Figure 8: **Synaptic architecture and behavioral contributions of vestibulospinal neurons.** Circuitry: Vestibulospinal cells receive convergent high amplitude excitatory inputs (green) from irregular afferents originating with the ipsilateral utricle (see also [32]), low-amplitude excitatory inputs (blue) from extra-vestibular sources and inhibitory inputs (red) from either the ipsilateral or contralateral utricle. Behavior: Vestibulospinal cells are involved with three key computations. The first two, movement timing and corrective kinematics together allow fish to maintain postural stability. The third allow fish to coordinate fin and body rotations to climb in the water column.

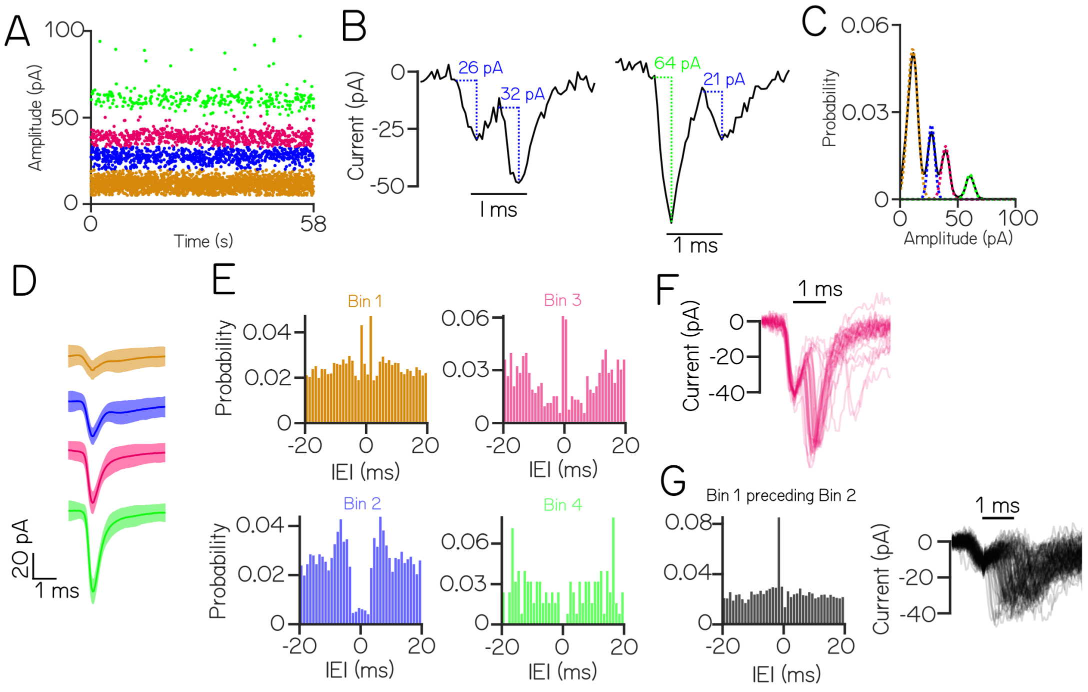


Figure S1: **Low-amplitude EPSCs events within the same amplitude bin reflect multiple neuronal inputs.** (A) An example cell with four stationary and discrete amplitude bins (color) (B) Example EPSC traces demonstrate events that co-occur within 1 ms. Event pairs are either within-bin (left) or across-bin (right) (C) To estimate an upper limit on the expected refractory period violations due to bin overlap, EPSC amplitude distributions were modeled as a sum of individual Gaussians (colors) (D) Average waveforms from each EPSC bin (± S.D.) (E) Autocorrelograms show structure of inter-event intervals within an EPSC bin; note peaks near zero in bins 1&3, a non-zero valley for bin 2, and a true valley for the high-amplitude bin 4 (F) Waveforms from EPSC pairs within bin 3 with latencies <2ms (G) Left: cross-correlogram between bin 1 and bin 2, Right: waveforms of EPSC pairs between 1 & 2 with latency <2ms. The small preceding event and large jitter between peaks are both inconsistent with the expected profile of an electrochemical synapse

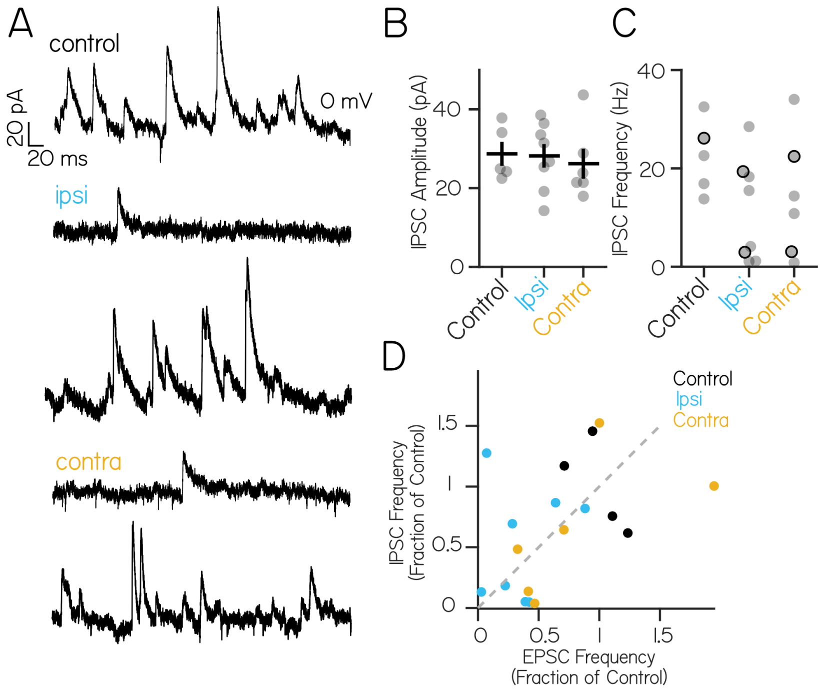


Figure S2: **Inhibitory currents onto a single vestibulospinal neuron can sometimes be disrupted by contralateral or ipsilateral utricular lesions; this disruption is not correlated with impairments to excitatory currents.** (A) Control trace from a neuron held at 0mV shows inhibitory input at rest (top). After ipsilateral (middle) or contralateral (bottom) otic capsule lesion, some cells experience strong loss of inhibitory currents, while others appear unaffected. (B) IPSC amplitude is unchanged across control and lesion conditions (Kruskal-Wallis H(2)=0.94, p=0.24) (C) The distribution of IPSC frequency after ipsilateral or contralateral lesions is bimodal. Black outlined dots represent cells for example traces in (A). No control neuron (n=5) experienced IPSC frequency less than 10 Hz. After ipsilateral lesions, 4/8 neurons experienced IPSC frequencies less than 10 Hz. After contralateral lesion, 2/6 neurons received IPSCs below 10 Hz (D) Spontaneous EPSC frequency and spontaneous IPSC frequency in recorded neurons from control (black), ipsilateral (blue) and contralateral (yellow) lesion conditions. Each dot is a neuron where both spontaneous excitatory and inhibitory current traces were recorded. For comparison, frequencies were normalized to the mean IPSC or EPSC frequency of control cells. Gray dashes indicate the unity line

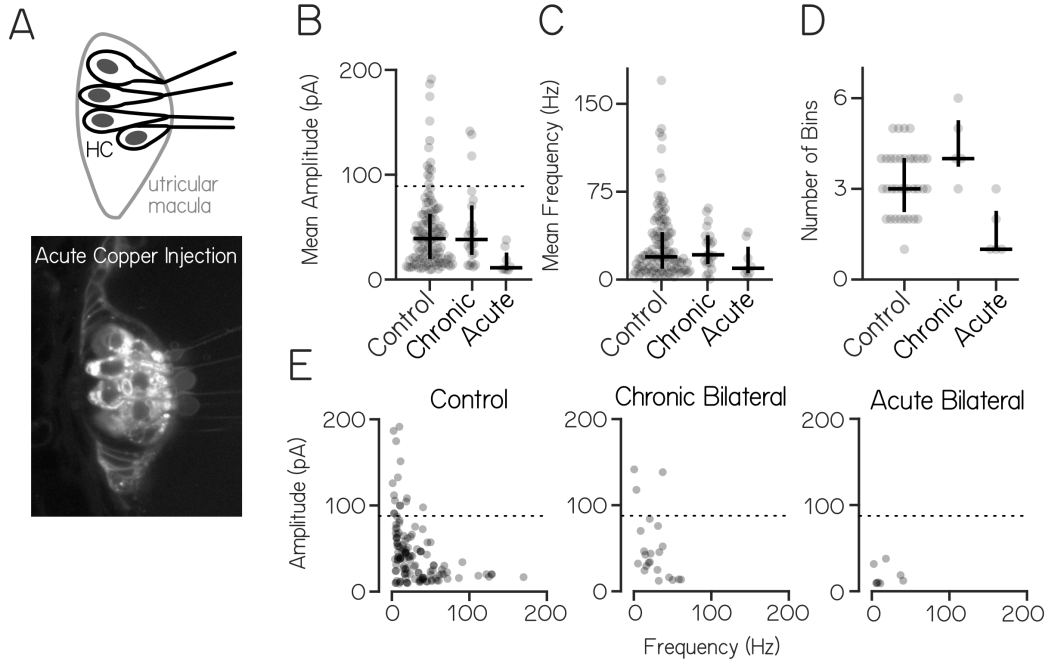


Figure S3: **Phasic tuning of action potentials, membrane potential, and synaptic events during linear translation.** (A) Phase of peak firing response to lateral and fore-aft stimuli. The majority of lateral-responsive cells respond to contralateral translation (6/10), and the majority of fore-aft responsive cells responded to rostral translation (7/10) (B) Representative low-pass filtered voltage traces during lateral translation from control (black), chronic (purple), and acute lesion (orange) shows rhythmic oscillation (solid) and sinusoidal fits (dashed) of membrane potential which are attenuated after lesion (C) During lateral translation, slow frequency voltage modulation is high in cells from control fish (3.4 mV, ± 0.5 SEM lateral, n=17 cells). Amplitude modulation is reduced after chronic (0.2 mV ± 0.2 SEM lateral, n=4 cells; p=0.02) but not acute utricle lesions (1. mV ± 0.7 SEM lateral, n=4 cells; p=0.16) (D) Slow frequency voltage modulation amplitude is positively correlated with the modulation in the mean EPSC event rate in both lateral (Pearson's r=0.59 ($R^2$=0.36) and fore-aft (Pearson's r=0.67 ($R^2$=0.45) translation (E-F) IPSCs and EPSCs from two cells show out-of-phase (Cell 1) or similar (Cell 2) responses to lateral translation. Top: acceleration with timing of peak response peak overlaid (blue/pink dashed line). Middle: spike rasters bottom: PSTH (F) Inhibitory and excitatory inputs from another example cell are co-tuned to lateral acceleration, with peak IPSC and EPSC rates occurring in-phase with ipsilateral acceleration G) Phase comparison of EPSC and IPSC peak rate of two example cells.

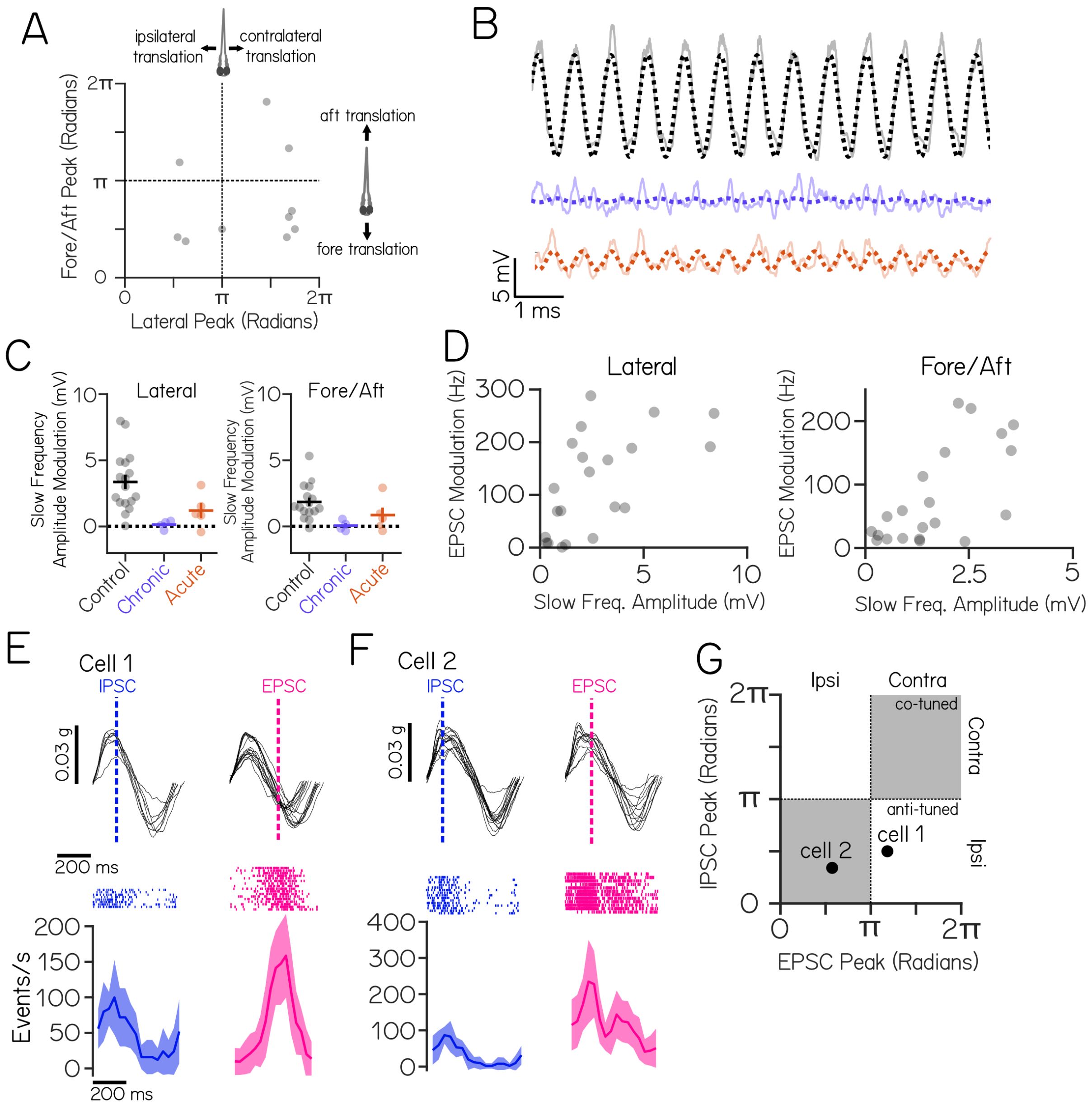


Figure S4: **Spontaneous excitatory synaptic input persists after chronic bilateral utricle loss (*otogelin* mutant) but is nearly abolished after acute bilateral lesions (chemical lesion of hair cells).** (A) Top: schematic of hair cells within the utricular macula of the inner ear. Bottom: disrupted hair cell membranes after copper injection (B) Mean amplitude of EPSC bins (gray circles) from vestibulospinal neurons in control, chronic and acute conditions. Median and inter-quartile range of data. "High amplitude" bins (top 5% of control) above black dashed line (C) Mean event frequency of EPSC bins. Median and inter-quartile range of data (D) Number of EPSC bins in a cell. Each dot is one vestibulospinal cell. Median and inter-quartile range of data (E) Mean amplitude of an EPSC bin (gray circle) as a function of frequency in control, chronic, and acute conditions. Dashed line delineates "high amplitude" bins

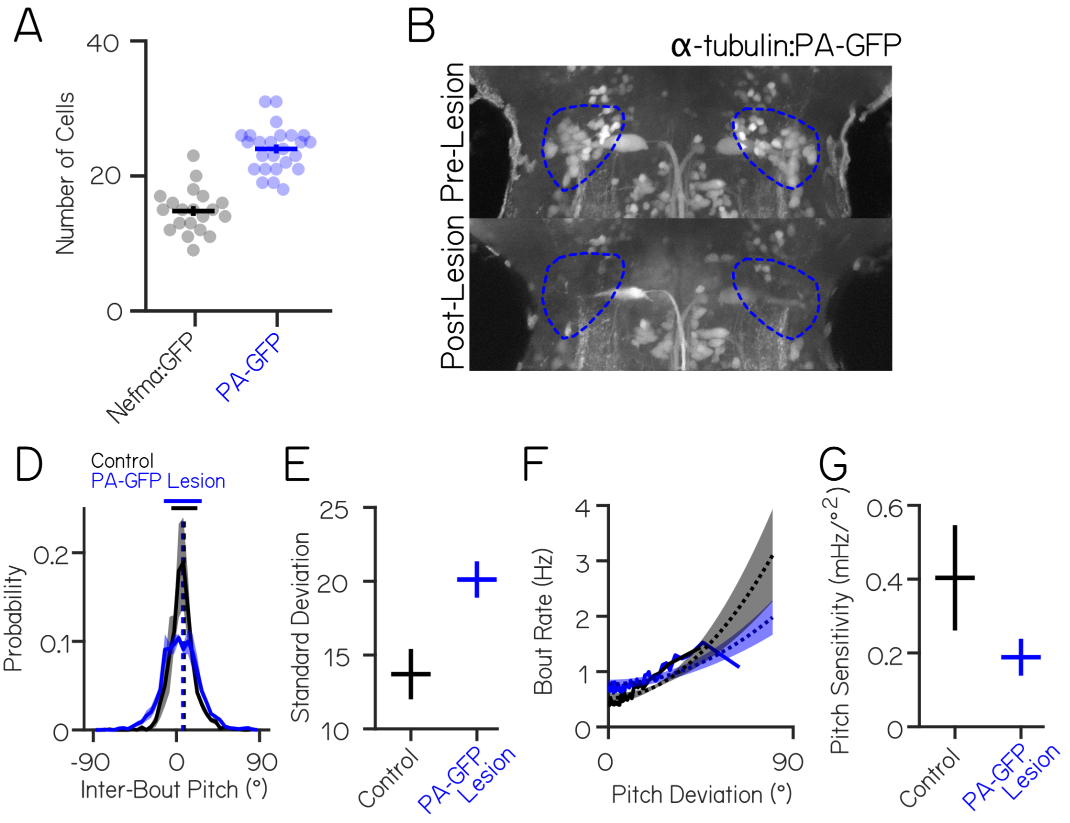


Figure S5: **Lesions of a larger pool of optically identified vestibulospinal neurons replicates postural and coordination disruption observed in *Tg(nefma:GAL4);Tg(UAS:GFP)* lesions.** (A) *Tg(nefma:GAL4);Tg(UAS:GFP)* labels fewer (14.8 ± 0.7 SEM, n=20 hemispheres, 10 fish) neurons than optically-backfilled *Tg(α-tubulin:C3PA-GFP)* fish (24 (± 0.6 SEM, n=25 hemispheres, 13 fish, unpaired t-test $p=8.3*10^{-12}$) (B) Dorsoventral distribution of vestibulospinal cells in both lines. *Tg(nefma:GAL4);Tg(UAS:GFP)* labels fewer cells in the dorsal vestibulospinal nucleus compared to photofills in *Tg(α-tubulin:C3PA-GFP)* (C) Representative maximum intensity projection of spinal projecting neurons in the hindbrain of a *Tg(α-tubulin:C3PA-GFP)* fish following optical backfill before (top) and after (bottom) two-photon mediated photoablation (D) Average probability distributions (± S.D) of inter-bout pitch angle for sibling controls (black, n=17 fish, 1828 bouts control) and vestibulospinal lesioned fish (blue, n=17 fish, 2125 bouts lesion) show no change in average posture (dashed vertical lines) but greater variability (solid horizontal lines ± 1 S.D.) (E) Standard deviation of pitch is higher in vestibulospinal lesioned fish (20.1±1.2) compared to sibling controls (13.7 ±1.7, paired t-test $p=1.0*10^{-5}$) (F) Bout rate as a function of deviation from preferred posture for lesions (blue) and control siblings (black). Solid lines represent raw data, dashed lines represent parabolic fits to raw data ± S.D. of the fit estimates. (G) Pitch sensitivity (parabolic fit) is decreased in vestibulospinal lesioned fish (0.2 ± 0.05 S.D., paired t-test p=0.002) compared to sibling controls (0.4 ± 0.14 S.D.)
